# Ki-67 shapes the nucleolus by anchoring chromatin via its amphiphilic properties

**DOI:** 10.1101/2025.07.23.666339

**Authors:** Daja Schichler, Yuki Hayashi, Letitia Fernandez, Mariam Chupanova, Alberto Hernandez-Armendariz, Beate Neumann, Sara Cuylen-Haering

**Affiliations:** Cell Biology and Biophysics Unit, European Molecular Biology Laboratory (EMBL), Heidelberg, Germany; Collaboration for Joint PhD Degree between EMBL and Heidelberg University, Faculty of Biosciences, Heidelberg, Germany; Advanced Light Microscopy Facility, European Molecular Biology Laboratory (EMBL), Heidelberg, Germany; Discovery Sciences, R&D, AstraZeneca, Gothenburg, Sweden; Max Planck Institute of Molecular Cell Biology and Genetics, Dresden, Germany; Cluster of Excellence Physics of Life, TU Dresden, Dresden, Germany

## Abstract

The nucleolus, a membrane-less organelle essential for ribosome biogenesis, adopts variable shapes across cell types and in response to environmental conditions, yet the mechanisms regulating its morphology and functional implications remain unclear. Using a high-throughput screen, we identify the proliferation marker Ki-67 as a central regulator of nucleolar shape. Ki-67 localises to the chromatin-nucleolus interface, where its depletion induces nucleolar rounding and reduces chromatin enrichment within and around the nucleolus. This effect is driven by amphiphilic properties conferred by two distinct affinity domains separated by a spacer. Given that chromatin loss is a common feature of rounded nucleoli in our screen, and acute chromatin digestion also induces rounding, we propose that the chromatin environment in and around the nucleolus plays a key role in determining nucleolar shape. Our study elucidates a novel Ki-67-mediated chromatin anchoring mechanism, tightly linking nucleolar shape to genome organisation and expanding our understanding of condensate morphology.

## Introduction

Membrane-less compartments are essential for cellular organisation and participate in diverse cellular functions. Unlike membrane-bound organelles, these compartments are dynamic and often exhibit liquid-like properties, allowing them to react quickly to stimuli. In both cells and in vitro, membraneless compartments typically adopt spherical shapes^1–3^, driven by surface tension to minimise their surface area^4^. However, some membrane-less compartments, such as nucleoli or nuclear speckles, commonly display irregular shapes^5^. This irregularity may be influenced by several factors such as their internal molecular properties, their interactions with the surrounding environment, or their viscoelastic properties. The precise molecular mechanisms and factors that determine the shape of specific membrane-less compartments remain, however, largely unknown.

Nucleoli are the largest membrane-less organelles in the nucleus, coordinating the process of ribosome biogenesis. To achieve this, they are organised into three distinct, nested subcompartments: the innermost fibrillar centre (FC), surrounded by the dense fibrillar component (DFC), and the outermost granular component (GC), which envelops dozens of FC-DFC subcompartments. This layered subcompartmentalisation is critical for spatially segregating different steps of ribosome biogenesis^6^, ensuring that transcription, processing, and assembly occur efficiently within specialised regions of the nucleolus.

In addition, nucleoli play a key role in the cellular stress response. Under stress conditions, such as DNA damage or heat shock, nucleoli undergo dynamic changes to adapt to the altered cellular environment. For instance, DNA damage can lead to the inhibition of ribosome biogenesis through the suppression of RNA polymerase I activity. This triggers the rounding of nucleoli and the reorganisation of their nested structure, resulting in the appearance of FC-DFC subcompartments on the surface of the GC region, a structure known as nucleolar caps^7^.

Another important function of nucleoli is their role in chromatin organisation. Nucleoli not only organise the ribosomal RNA (rRNA) gene clusters, but they are also surrounded by a rim of dense heterochromatin^8^. These heterochromatic regions, known as nucleolar-associated domains (NADs)^9^, are enriched in telomeres and other repetitive DNA sequences and regions with low gene density^10^. Therefore, nucleoli are considered to be one of the central structures for the organisation of inactive heterochromatin, contributing to the spatial arrangement and regulation of these genomic regions.

Despite the liquid-like properties of nucleoli^11–13^, their shape is highly variable between cell types^14^. While some cell types exhibit nearly spherical nucleoli, others display highly irregular morphologies, characterised by alternating concave and convex regions. Such shape diversity is not only a feature of normal cellular differentiation but also a hallmark of certain diseases. In conditions like viral infections^15,16^, neurodegeneration^17^, and cancer^18^, changes in nucleolar shape, size, and number have been observed and are often used as clinical diagnostic markers^19–21^, suggesting that nucleolar shape is tightly regulated and responsive to the cellular environment. These alterations in shape may reflect underlying changes in ribosome biogenesis, stress signalling, or genome stability. However, the molecular mechanisms driving these shape changes remain poorly elucidated.

Here, we report the results of an RNA interference (RNAi) screen to identify proteins involved in regulating nucleolar shape. This screen identified several candidate proteins, including the cell proliferation marker Ki-67, that induce nucleolar rounding upon their depletion. In addition to inducing nucleolar rounding, Ki-67 depletion led to the loss of chromatin from both the nucleolar rim and its interior. Quantitative live imaging and protein engineering experiments revealed that Ki-67 localises at the chromatin-nucleolar interface, dependent on its two distinct affinity domains. This interaction promotes chromatin enrichment and the maintenance of irregular nucleolar shape. Notably, loss of chromatin enrichment in the nucleolus was observed in most of the candidate proteins from our RNAi screen that induce nucleolar rounding. In most cases, Ki-67 expression levels were reduced, suggesting that Ki-67 may play an essential role in regulating chromatin distribution in the nucleolus. Since acute chromatin digestion was sufficient to induce nucleolar rounding, the chromatin network is most likely a central determinant of nucleolar morphology. Our findings highlight the role of chromatin in regulating nucleolar shape, with Ki-67-dependent chromatin recruitment facilitated by its amphiphilic structure emerging as one of the central molecular mechanisms underlying the irregular morphology of nucleoli.

## Results

### Identification of nucleolar shape regulators

To identify molecular factors regulating nucleolar shape, we conducted a live-cell imaging-based RNAi screen in a HeLa cell line expressing NPM1-EGFP as a nucleolus marker and H2B-mCherry as a chromatin marker. We targeted 614 genes using 2–3 individual siRNAs per gene, focusing on a library enriched for nucleolus-associated proteins (see Table S1), including proteins that regulate nucleolar number^22^ and are associated with phase separation ability^23,24^. Live cells were imaged using an automated wide-field microscope, and their cell-cycle stages were classified through supervised machine learning. We extracted interphase cells and quantified nucleolar aspect ratio as a proxy for shape (Figure 1A). While nucleoli in cells treated with a non-targeting control siRNA were typically irregularly shaped, our screen identified many candidates that induce nucleolar rounding (Figure 1B and 1C).

**Figure 1.**
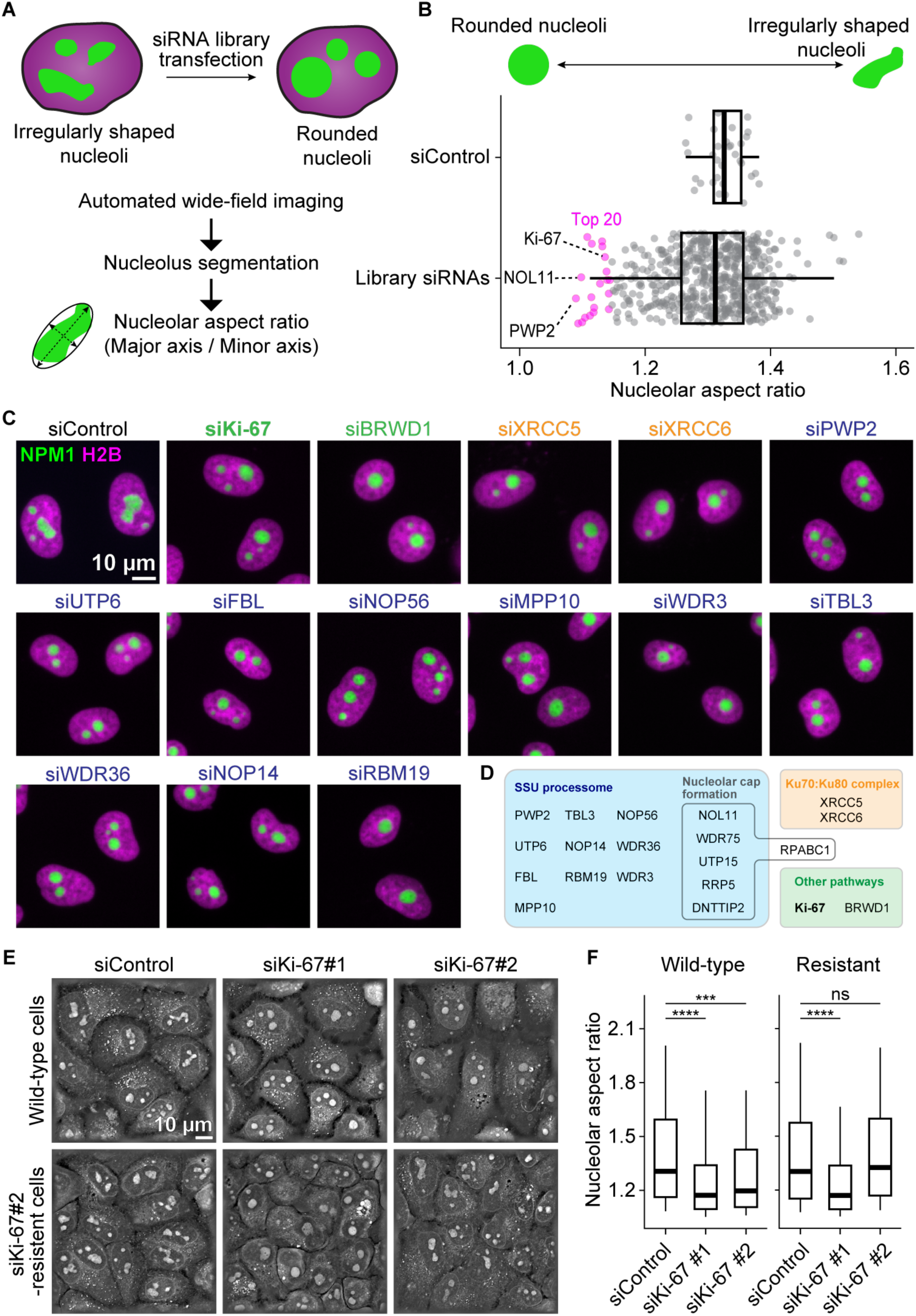
An RNAi screen for genes regulating nucleolar shape identifies Ki-67. (A) Experimental design of a live cell RNAi screen to identify genes regulating nucleolar shape. HeLa cells expressing H2B-mCherry (magenta) and NPM1-EGFP (green) were seeded in multiwell plates with siRNAs targeting one gene per well and imaged after 72 h. Nucleoli were segmented using an automated image analysis pipeline, and their shape was quantified by their aspect ratio after fitting an ellipse around each nucleolus. (B) Outcome of the RNAi screen targeting 614 genes. Individual data points in a non-targeting control siRNA (siControl) indicate median nucleolar aspect ratio per well, providing a baseline for comparison. Data points from the library siRNAs indicate median values per gene based on 2 or 3 different siRNAs. Magenta-coloured data points highlight the top 20 genes inducing nucleolar rounding. Boxplots display the median (centre line), interquartile range (box), and whiskers extending to the 10th and 90th percentiles. (C, D) Raw data from the screen showing the effect of knocking down the top 20 genes identified in (B). Chromatin is labelled by H2B-mCherry (magenta), and nucleoli are marked by NPM1-EGFP (green). The molecular function of the top 20 genes has been manually annotated and classified into different colour-coded groups. (E) Label-free holotomographic live imaging of HeLa cells 72 h after transfection. Wild-type HeLa cells (top panels) or HeLa cells resistant to siRNA Ki-67#2 (bottom panels) were transfected with siControl or two different Ki-67 siRNAs (siKi-67#1 and siKi-67#2). A single z-slice is shown. (F) Quantification of nucleolar aspect ratio in holotomographic images. The aspect ratio of nucleoli was measured for wild-type HeLa cells (left plots) or HeLa cells resistant to siKi-67#2 (right plots). Boxplots display the median (centre line), interquartile range (box), and whiskers extending to the 10th and 90th percentiles. For (F), n = 299 nucleoli (siControl, wild-type), 194 nucleoli (siKi-67#1, wild-type), 238 nucleoli (siKi-67#2, wild-type), n = 390 nucleoli (siControl, resistant), 309 nucleoli (siKi-67#1, resistant), 343 nucleoli (siKi-67#2, resistant), 2 experiments. ****P<0.0001 and ***P=0.004, a Kolmogorov-Smirnov test. See also Figure S1 and S2.

Remarkably, 15 of the 20 highest-scoring candidates were components of the small subunit (SSU) processome (Figure 1C and 1D), crucial for ribosome biogenesis^25^. Other notable candidates included XRCC5 and XRCC6, a heterodimer involved in the DNA double-strand break repair. Additionally, depletion of RPABC1 (encoded by the POLR2E gene), a subunit common to all three RNA polymerases, the Bromodomain and WD repeat-containing protein 1 (encoded by BRWD1)^26^, and the well-known proliferation marker Ki-67^27,28^ (encoded by MKI67) also induced nucleolar rounding.

Since knockdown of SSU processome components impairs rRNA processing^29^, the observed nucleolar rounding might be a consequence of stalled ribosome biogenesis and subsequent reorganisation^30^. To test this hypothesis, we performed a secondary screen using a dual colour cell line labelling inner and outer nucleolar subcompartments (Figure S1). As expected, knockdown of the RNA polymerase subunit RPABC1 led to nucleolar reorganisation with a characteristic cap-like structure. Similarly, knockdown of multiple SSU processome components induced nucleolar cap formation, suggesting that rounding is likely an indirect consequence of stalled ribosome biogenesis. While XRCC5 and XRCC6 depletion did not induce caps under our conditions, they likely affect nucleoli structure indirectly through genomic instability^31^. BRWD1 is thought to function as a transcription regulator that is involved in regulating cell morphology^32^. Therefore, its effect might be indirect, through the regulation of other genes. Ki-67, on the other hand, localises to nucleoli during interphase^33^ and is dispensable for efficient rDNA transcription and pre-rRNA processing^34^, suggesting that nucleolar rounding must occur via a different pathway. Notably, Ki-67 has been proposed to function as a surfactant on the chromosome surface during mitosis^35^. Although it remains unclear whether its surfactant function persists in interphase^36,37^, Ki-67’s reported localisation and suggested function make it a compelling candidate for regulating nucleolar morphology during interphase. Based on this rationale, we investigated Ki-67’s role in shaping nucleoli.

To avoid artefacts caused by overexpression of nucleolar marker proteins or fluorescent protein tags, we performed label-free holotomographic imaging and used supervised machine learning to segment nucleoli^38^ (Figure S2A). To determine whether nucleolar rounding was specific to Ki-67 depletion rather than a non-specific siRNA effect, we transfected two different Ki-67 siRNAs, both of which deplete Ki-67 to background levels (Figure S2B and S2C), into wild-type HeLa cells and a cell line resistant to one of the siRNAs due to Cas9-engineered synonymous mutations of the target region^35^. In wild-type cells, Ki-67 depletion led to rounder nucleoli, while control cells had irregularly shaped nucleoli (Figure 1E, top row, and 1F). In the resistant cell line, only the siRNA targeting the non-mutated region caused nucleolar rounding, whereas the siRNA to which the cells were resistant resulted in irregular nucleoli, resembling the control (Figure 1E, bottom row, and 1F). The nucleolar rounding phenotype was also observed in unlabelled Ki-67 knockout (KO) cells (Figure S2D and S2E). In summary, these results confirmed that the nucleolar rounding induced by transfection with Ki-67-targeting siRNAs was a specific effect of Ki-67 knockdown.

### Ki-67 localises to the interface of chromatin and the nucleolus

To understand how Ki-67 regulates nucleolar shape, we revisited its localisation within nucleoli. Several studies have described its localisation to the region surrounding nucleoli, known as the nucleolar periphery or rim^33,34,39^. Additionally, early studies using immunoelectron microscopy or immunofluorescence have demonstrated that Ki-67 also localises to the inner regions of nucleoli with limited overlap with the other nucleolar subcompartments^39–41^. However, its exact localisation within these inner nucleolar regions remained unclear.

To investigate the precise localisation of Ki-67 in nucleoli, we generated a HeLa cell line endogenously tagged with EGFP at the Ki-67 N-terminus and stably expressing the nucleolar protein NPM1 tagged with a SNAP-tag. We then performed live-cell Airyscan super-resolution imaging of Ki-67 and NPM1, along with DNA staining. Although Ki-67 was enriched at the nucleolar rim, it was also clearly detectable within nucleoli (Figure 2A–2D). This internal Ki-67 signal did not overlap with the canonical nucleolar subcompartments (Figure S2), consistent with previous protein localisation studies^39,40^. Since we also observed a significant chromatin signal in the nucleolar interior (Figure 2A and 2E), we tested for co-localisation with Ki-67. Line profile analyses along intra-nucleolar chromatin foci indicated that Ki-67 signal peaks were positioned between DNA and NPM1 signal peaks (Figure 2F, line 1). Similarly, at the nucleolar rim, Ki-67 signal was most intense between the NPM1 and DNA signal peaks (Figure 2F, line 2). These results suggest that Ki-67 localises at the boundary between the nucleolus (GC compartment) and chromatin.

**Figure 2.**
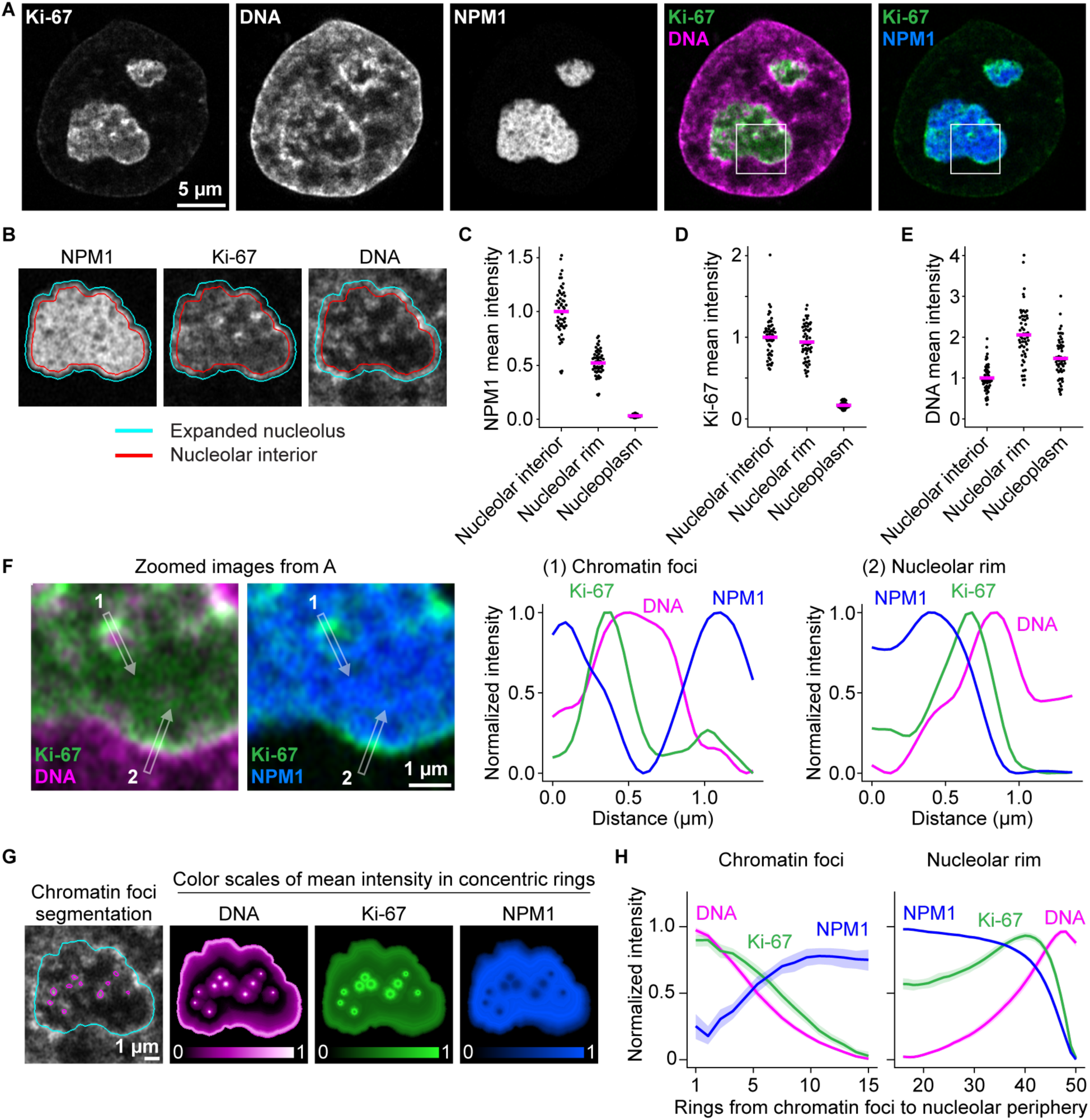
Ki-67 localises at the chromatin-nucleolus interface. (A) Live cell Airyscan imaging of HeLa cells. Cells endogenously tagged with EGFP-Ki-67 and stably expressing SNAP-NPM1 were stained with SPY555-DNA dye. SNAP-NPM1 was labelled with SNAP-silicon rhodamine (SiR). (B) Nucleolar segmentations based on the NPM1 signal. The nucleolar segmentation was expanded (cyan) and shrunk (red; nucleolar interior) by 5 pixels, corresponding to approximately 200 nm. The nucleolar rim was defined by subtracting the shrunk segmentation from the expanded segmentation. (C–E) Quantification of NPM1 (C), Ki-67 (D), and DNA (E) signal intensities in the nucleolar interior, rim and nucleoplasm. Mean intensities in each region were normalised to the mean intensity in the nucleolar interior. Bars represent mean values. (F) Signal distribution of NPM1, Ki-67, and DNA at the intra-nucleolar chromatin foci and the nucleolar periphery. Images show the regions indicated by white squares in (A). Line profile measurements were performed around chromatin foci (1) and the nucleolar rim (2). Signal intensities were normalised to the respective maximum and minimum fluorescence intensities. (G) Quantification of signal distributions of Ki-67, DNA, and NPM1 from the intra-nucleolar chromatin foci. Segmentations of intra-nucleolar chromatin foci (magenta) and expanded nucleolus (cyan) are highlighted in the leftmost image. The nucleolus was divided into 50 concentric rings from the centre of chromatin foci to the edge of the expanded nucleolus. The colour scale indicates relative signal intensity. (H) Quantification of the signal intensity in concentric rings. Mean intensities in each concentric ring were normalised by min-max scaling, applied independently to rings 1–15 (around the chromatin foci) and rings 16–50 (around the nucleolar rim). Line and shaded area indicate mean ± 95% confidence interval. For (E), (F) and (H), n = 53 nuclei, 3 experiments. See also Figure S3.

To quantitatively confirm this observation across many cells, we segmented intra-nucleolar chromatin and divided the nucleolus into 50 concentric rings around the chromatin foci’s centre points (Figure 2G). We then measured Ki-67, DNA, and NPM1 signal intensities in each ring, enabling us to generate spatial intensity profiles across multiple cells (Figure 2H). Adjacent to the chromatin foci (rings 1–15), Ki-67 displayed a broader peak compared to DNA, suggesting its localisation on the surface of intra-nucleolar chromatin foci. At the nucleolar boundary (rings 15–50), Ki-67 intensity peaked concomitantly with a sharp decrease in NPM1 signal, preceding the peak in DNA signal. These results corroborate our line profile analyses, reinforcing the conclusion that Ki-67 preferentially localises at the boundary between chromatin and the nucleolus, both at the nucleolar rim and within nucleoli.

### Ki-67 anchors chromatin into the nucleolar interior

Given that Ki-67 localises at the chromatin-nucleolus boundary, we next investigated whether Ki-67 plays a role in regulating the chromatin environment within and around nucleoli. To test this, we depleted Ki-67 using siRNAs in HeLa cells stably expressing histone H2B-mCherry as a chromatin marker and NPM1-EGFP as a nucleolus marker (Figure 3A–3E). Depletion of Ki-67 resulted in the formation of rounder nucleoli as expected, but had only a minor impact on nucleolar size (Figure 3B and 3C). In addition, Ki-67 depletion not only decreased the H2B signal intensities in the nucleolar rim (Figure 3E), in line with previous reports^34,42^, but also led to reduced H2B signal intensities in the nucleolar interior (Figure 3D), causing the nucleoli to appear as dark voids (Figure 3A). We conclude that Ki-67 is important for maintaining chromatin association not only at the periphery but also in the interior of nucleoli.

**Figure 3.**
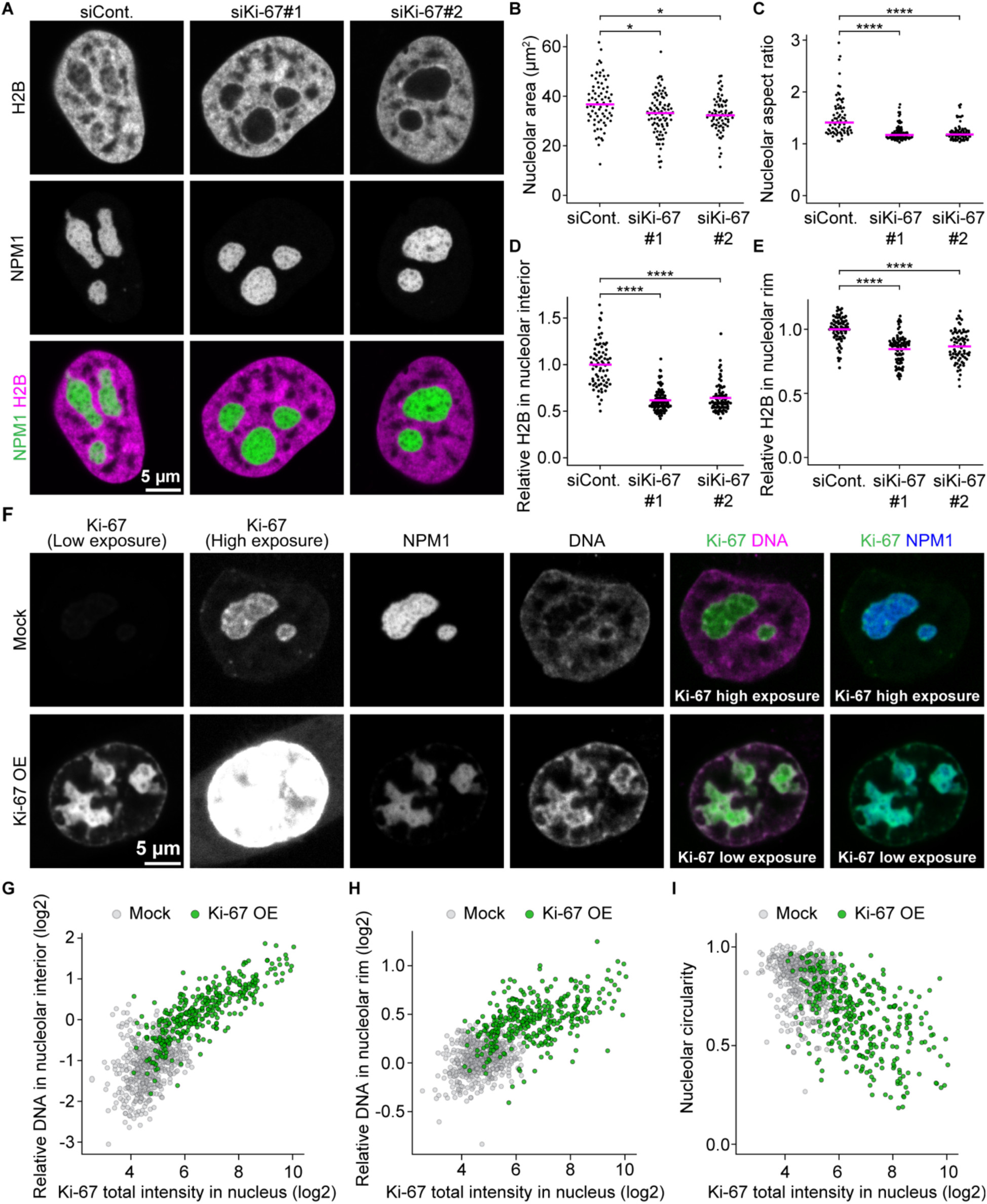
Ki-67 regulates chromatin loading into nucleoli. (A) Live imaging of HeLa cells stably expressing H2B-mCherry and NPM1-EGFP. Cells were transfected with a non-targeting control siRNA (siCont.) or two different Ki-67 siRNAs, followed by confocal imaging 72 h post-transfection. A single z-slice is shown. (B, C) Quantification of nucleolar size and shape from NPM1 signals. Nucleoli were segmented based on NPM1 signals to measure total nucleolar area per nucleus (B) and median nucleolar aspect ratio per nucleus (C). Bars indicate median values. (D, E) Quantification of chromatin enrichment in the nucleolar interior and its rim. Relative H2B signal intensities were calculated by dividing mean intensities in the nucleolar interior (D) or the nucleolar rim (E) by mean intensities in the nucleoplasm. Bars indicate mean values. (F) Live imaging of Ki-67 overexpressing cells (Ki-67 OE). Cells endogenously tagged with EGFP-Ki-67 and stably overexpressing SNAP-NPM1 were transfected with an EGFP-Ki-67 plasmid and isolated by FACS. SNAP-NPM1 was labelled with SNAP-SiR, and DNA was stained with SPY555-DNA. Two different Ki-67 signal intensities are shown by adjusting brightness and contrast settings. Mock refers to cells treated with the transfection reagent only. (G, H) Ki-67 expression level-dependent chromatin enrichment in the nucleolar interior (G) and its rim (H). Relative DNA signal intensities, calculated as described for the H2B signal intensities in (D, E), are plotted against the total EGFP-Ki-67 intensity in the nucleus. (I) Ki-67 expression level-dependent increase in the irregularity of nucleolar shape. Median circularity of segmented nucleoli from NPM1 signals per nucleus is plotted against the total intensity of EGFP-Ki-67 in the nucleus. For (B–E), n = 72 nuclei (siControl), 87 nuclei (siKi-67#1), 75 nuclei (siKi-67#2), 2 experiments. **** P < 0.0001 and * P < 0.05 with Kruskal–Wallis test followed by Dunn’s test, compared to siControl. For (G–I), n = 517 nuclei (Mock); n = 355 nuclei (Ki-67 OE), 2 experiments. See also Figure S4 and S5.

Based on this finding and considering the localisation of chromatin and Ki-67 in the nucleolar interior between nucleolar subcompartments, we investigated whether chromatin tethering by Ki-67 could support the separation of nucleolar subcompartments. Interestingly, Ki-67 depletion did not affect the internal organisation of the nucleolar subcompartments, as both the number and area of UBF foci remained constant (Figure S4), consistent with the reports that Ki-67 is not required for ribosome biogenesis^34^. These findings indicate that the separation of internal nucleolar subcompartments does not depend on Ki-67’s chromatin recruitment to the nucleolar interior.

To further investigate Ki-67’s role in anchoring chromatin to the nucleolus, we next tested whether Ki-67 overexpression causes additional chromatin enrichment within and around nucleoli. We transiently overexpressed EGFP-Ki-67 in cells that already expressed endogenous EGFP-tagged Ki-67 and stably expressed SNAP-NPM1. After FACS-sorting to isolate cells with higher Ki-67 expression compared to the endogenous levels (Figure 3F and S5A), we quantified the mean intensities of DNA and Ki-67 in the nucleolar interior and the rim (Figure 3G and 3H). The DNA signal intensities in the nucleolar interior and the rim increased with EGFP-Ki-67 expression (Figure 3F–3H). At very high levels, Ki-67 recruited substantial amounts of chromatin to the nucleolar interior, resulting in low chromatin densities in the rest of the nucleus (Figure 3F). Analysis of DNA and NPM1 signals in concentric rings from the nucleolar centre to the periphery confirmed this observation (Figure S5B and S5C): While the DNA enriched at the rim in both mock and Ki-67 overexpression conditions, the normalised peak observed for Ki-67 overexpression was broader and extended into the nucleolar interior, possibly due to saturation at the rim. Together, these results demonstrate that Ki-67 overexpression enhances chromatin recruitment to the nucleolar interior and the rim.

If chromatin incorporation contributes to nucleolar shape, an increase in Ki-67 expression might lead to higher nucleolar irregularity. Indeed, cells with high Ki-67 expression displayed highly irregular nucleoli (Figure 3F). However, the nucleolar aspect ratio did not correlate with Ki-67 expression (Figure S5D), likely due to its reliance on ellipse fitting (Figure 1A), which underestimates nucleolar irregularity. In contrast, nucleolar circularity, which considers perimeter and area, captured nucleolar irregularity much better and revealed a clear negative correlation with Ki-67 levels (Figure 3I), indicating that nucleolar irregularity depends on Ki-67 expression. Overall, these results demonstrate that Ki-67 recruits chromatin to the nucleolar interior and the rim and modulates nucleolar shape in a dose-dependent manner.

### Nucleolar rounding is reversible and a direct consequence of Ki-67 removal

Since the siRNA-mediated protein knockdown generally takes 2–3 days to achieve efficient protein depletion, it was unclear whether the observed nucleolar rounding and chromatin removal from nucleoli was a direct consequence of Ki-67 depletion or resulted from secondary effects such as alterations in gene expression by altering chromatin organisation^34^. To test this, we employed an auxin-inducible degron system^43,44^ for the rapid degradation of Ki-67. We generated a knock-in cell line endogenously expressing Ki-67 tagged with EGFP and a miniDegron-tag^45^ at its N-terminus and stably expressing the F-box protein OsTIR1 and the nucleolar marker FBL-TagRFP. Treatment with indole-3-acetic acid (IAA) rapidly decreased Ki-67 fluorescence to background levels within 2 h (Figure 4A and 4B). Removal of IAA from the culture medium restored Ki-67 fluorescence levels to approximately 90% at 14 h after washout (Figure 4B and 4C, last timepoint), providing an opportunity to assess the correlation between acute changes in Ki-67 levels and nucleolar rounding and intra-nucleolar chromatin levels.

**Figure 4.**
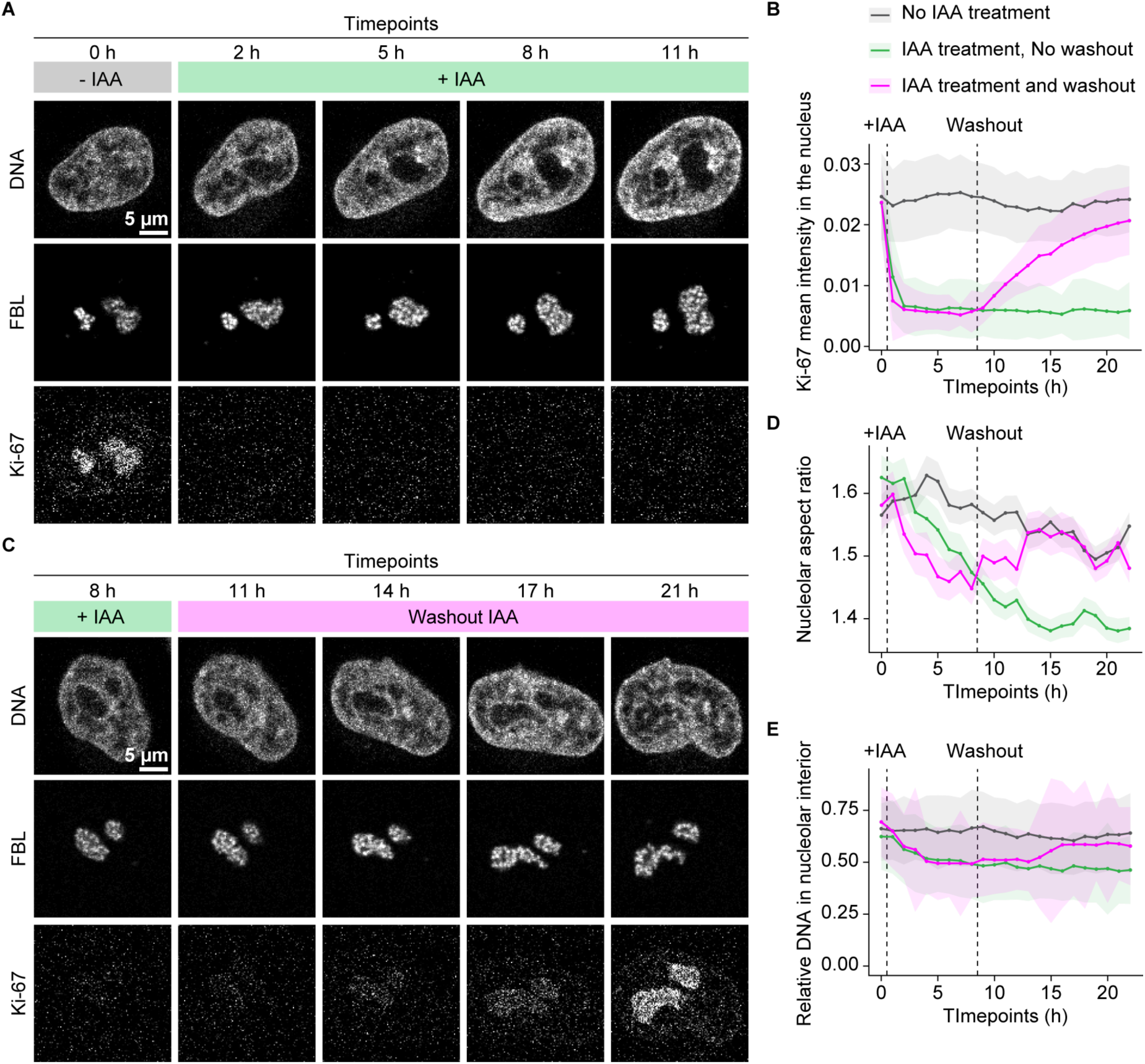
Nucleolar rounding and chromatin enrichment in the nucleolus are reversible and direct consequences of Ki-67 removal. (A) Acute degradation of Ki-67 and its effect on nucleolar shape and chromatin enrichment in the nucleolus. HeLa cells endogenously expressing EGFP-AID-Ki-67 and stably overexpressing FBL-TagRFP cells were treated with 3-Indole-acetic acid (IAA) 0.5 h after the start of time-lapse imaging. DNA was stained with SiR-DNA. Single z-slice of a representative example quantified in (C–E) is shown. (B) Recovery of Ki-67 expression after IAA washout and its effect on the nucleolar shape and chromatin enrichment in the nucleolus. IAA treatment was performed as in (A) and washed out 8.5 h after the start of time-lapse imaging. (C–E) Quantification of Ki-67 signal decay and recovery (C), nucleolar aspect ratio (D), and relative DNA signal intensities in the nucleolus over the nucleus (E) of indicated conditions using images as in and (B). For (C–E), n = 193 to 391 nuclei (No IAA treatment), n = 171 to 381 nuclei (IAA treatment, No washout), n = 161 to 298 nuclei (IAA treatment, Washout) per time point, 2 experiments. See also Figure S6.

The addition of IAA immediately decreased chromatin levels in the nucleolar interior and induced nucleolar rounding within 2–3 h. While intra-nucleolar chromatin levels reached their minimum after 5 h, rounding continued to decrease until 15 h post-treatment (Figure 4D and 4E). Importantly, the changes in intra-nucleolar chromatin levels and nucleolar roundness began to recover after the IAA washout. Both parameters required approximately 7 h for recovery, by which time Ki-67 levels had reached about 54% of the pre-depletion levels. Neither parameter fully recovered to the initial baseline value within the time imaged, possibly due to incomplete recovery of EGFP-miniDegron-Ki-67 expression. As an IAA-insensitive control, we used a cell line expressing wild-type Ki-67 and FBL-TagRFP (Figure S6A). The nucleolar shape and intra-nucleolar chromatin of IAA-insensitive control cells were largely unaffected by imaging conditions with or without IAA treatment (Figure S6B and S6C). Importantly, the observed alterations of nucleolar shape and chromatin enrichment upon Ki-67 depletion and their recovery were evident in the absence of cell division (Figure 4A and 4C). Thus, the rapid changes in nucleolar roundness and chromatin enrichment following Ki-67 protein depletion and recovery provide strong evidence that Ki-67 directly regulates both nucleolar morphology and chromatin enrichment within the nucleolar interior.

### Ki-67’s amphiphilic properties are necessary for its localisation and function in interphase

To investigate how Ki-67 mediates nucleolar chromatin incorporation and concurrent shape changes, we tested which specific domain of Ki-67 is required for these functions. Ki-67 is a large (360 kDa) disordered protein that consists of mainly three regions (Figure 5A): a C-terminal leucine/arginine-rich region (LR domain) implicated in DNA binding^46,47^, a middle region characterized by 16 repeats (Repeats), and an N-terminal domain containing a highly positively charged patch (CP domain) involved in clustering of chromosomes during anaphase^37^. The N-terminus also includes two short structured domains: a forkhead-associated (FHA) domain implicated in phosphopeptide binding^27,48^ and a Protein phosphatase 1 (PP1) binding motif^49^.

**Figure 5.**
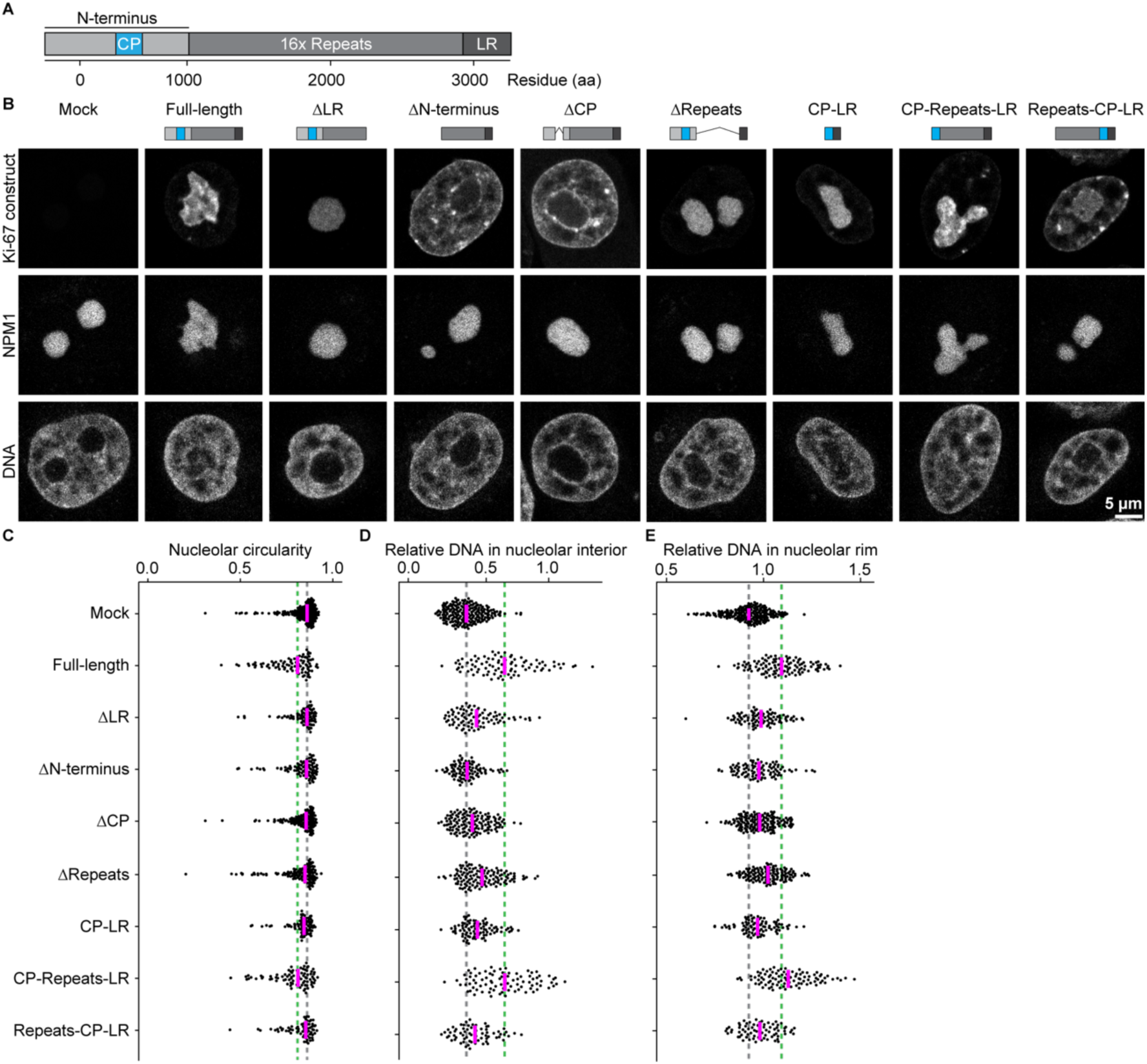
Ki-67’s dual affinity domains, separated by the repeat region, regulate the nucleolar roundness and chromatin enrichment in the nucleolus. (A) Schematic representation of Ki-67 domains. CP: positively charged patch of 186 amino acids, LR: leucine/arginine-rich DNA-binding domain. (B) Live imaging of Ki-67 KO HeLa cells transfected with Ki-67 domain mutants. Cells stably expressing mTurquoise2-NPM1 and transfected with EGFP-tagged Ki-67 mutant plasmids were collected by FACS and imaged on a confocal microscope. DNA was stained with SiR-DNA. (C) Quantification of nucleolar circularity in Ki-67 KO cells expressing different Ki-67 domain mutants. The nucleolus was segmented based on NPM1 signals, and median nucleolar circularity per nucleus was calculated for cells expressing Ki-67 mutants at levels comparable to endogenous Ki-67. Dashed lines indicate median values of full-length Ki-67 (green) and mock-transfected cells (grey). (D, E) Quantification of chromatin enrichment in the nucleolus of cells expressing different Ki-67 mutants. Relative DNA signal intensities were measured in the nucleolar interior (D) and rim (E), as described in (Figure 3D and 3E), for cells expressing Ki-67 mutants at endogenous Ki-67 levels. Dashed lines indicate median values of full-length Ki-67 (green) and mock-transfected cells (grey). For (C–E), n = 220 nuclei (mock), n = 87 nuclei (full-length), n = 88 nuclei (ΔLR), n = 94 nuclei (ΔN-terminus), n = 139 nuclei (ΔCP), n = 34 nuclei (LR), n = 128 nuclei (ΔRepeats), n = 87 nuclei (CP-LR), n = 86 nuclei (CP-Repeats-LR), n = 69 nuclei (Repeats-CP-LR), 3 experiments. See also Figure S7.

We transiently expressed wild-type EGFP-Ki-67 and different truncation mutants in Ki-67 KO cells stably expressing mTurquise2-NPM1 as a nucleolus marker (Figure 5B), and FACS-sorted them using the cell line with endogenous EGFP-Ki-67 as a reference (Fig. S7A). In our analysis, we only considered cells with similar expression levels to endogenous EGFP-Ki-67 (Figure S7B-S7E). As expected, mock-transfected Ki-67 KO cells exhibited rounded nucleoli lacking chromatin (Figure 5B, Mock), and full-length Ki-67 localised at the chromatin-nucleolus boundary, restoring the wild-type phenotype with irregular nucleolar shape and chromatin enrichment (Figure 5B–5E, Full-length). Similar to our previous experiments (Figure 3F–3I), higher expression levels of full-length Ki-67 above the wild-type levels decreased nucleolar circularity and increased DNA intensity within nucleoli in a dose-dependent manner (Figure S7B–S7E, Full-length).

Previous studies have indicated that the C-terminal LR domain is important for chromatin binding, while the N-terminus is required for nucleolus localisation^47,50^. In line with this, our Ki-67 mutant lacking the LR domain (ΔLR) localised homogeneously within the nucleolus, while a Ki-67 mutant lacking the N-terminus region (ΔN-terminus) localised to chromatin (Figure 5B). Given our previous finding that the CP-domain of Ki-67 is critical for its phase separation with RNA^37^, we hypothesised that this domain might anchor Ki-67 to the nucleolus. Indeed, only the removal of the CP domain (ΔCP) relocalised Ki-67 from nucleoli to chromatin. Importantly, in contrast to full-length Ki-67, none of the mislocalised mutants (ΔLR, ΔN-terminus, ΔCP) were able to restore the irregular shape of nucleoli or the enrichment of chromatin to the nucleolus (Figure 5C–5E). These findings demonstrated that the LR and CP domains are essential for Ki-67’s localisation at the boundary between chromatin and nucleoli and for the regulation of the nucleolar shape as well as chromatin enrichment.

To test whether CP and LR are sufficient for Ki-67 localization and function, we tested a construct lacking the middle repeat region (ΔRepeats) as well as a direct CP-LR fusion. Surprisingly, both constructs localised homogenously to nucleoli and could only partially restore the irregular shape and chromatin enrichment (Figure 5B–5E). These findings indicate that while the CP and LR play a crucial role, they are insufficient on their own to fully rescue the Ki-67 wild-type phenotype. In contrast, a minimal Ki-67 construct with the repeats between CP and LR (CP-Repeats-LR) was able to localise correctly and restore nucleolar irregularity and chromatin enrichment (Figure 5B–5E), suggesting that the repeat region is also required for proper localisation and function. Interestingly, however, a shuffled minimal Ki-67 mutant (Repeats-CP-LR) did not restore these parameters efficiently even at higher expression levels than endogenous Ki-67 (Figure S7B–S7E). It is thus tempting to speculate that proper spacing between the CP and LR regions is necessary for the full functionality of Ki-67.

Based on the hypothesis that Ki-67 requires a spacer between the nucleolus and chromatin binding domains, we wondered whether it adopts an extended conformation at the chromatin-nucleolus border. To test this, we expressed a Ki-67 version with EGFP at one end and mCherry at the other (Figure 6A and 6B), or both fluorescence tags at the Ki-67 N-terminus (Figure 6C). We acquired high-resolution confocal images of nucleoli and measured the fluorescence signal intensity of both tags along line profiles from the nucleolar periphery inward. By fitting Gaussian functions (Figure 6D–6F), we determined peak positions in both channels and calculated the displacement between them (Figure 6G), indicating molecular extension and orientation. For EGFP-Ki-67-mCherry, the displacement was 37.7 ± 35.2 nm, indicating that Ki-67’s N-terminus protrudes towards the nucleolar interior. The reverse construct (mCherry-Ki-67-EGFP) showed a displacement of -29.6 ± 38.4 nm, while the control construct (EGFP-mCherry-Ki-67) had only 1.2 ± 24.8 nm, confirming that the previous measurements reflected molecular extension. Interestingly, this extension is comparable to Ki-67’s extension on the chromosome surface during mitotic exit^36^.

**Figure 6.**
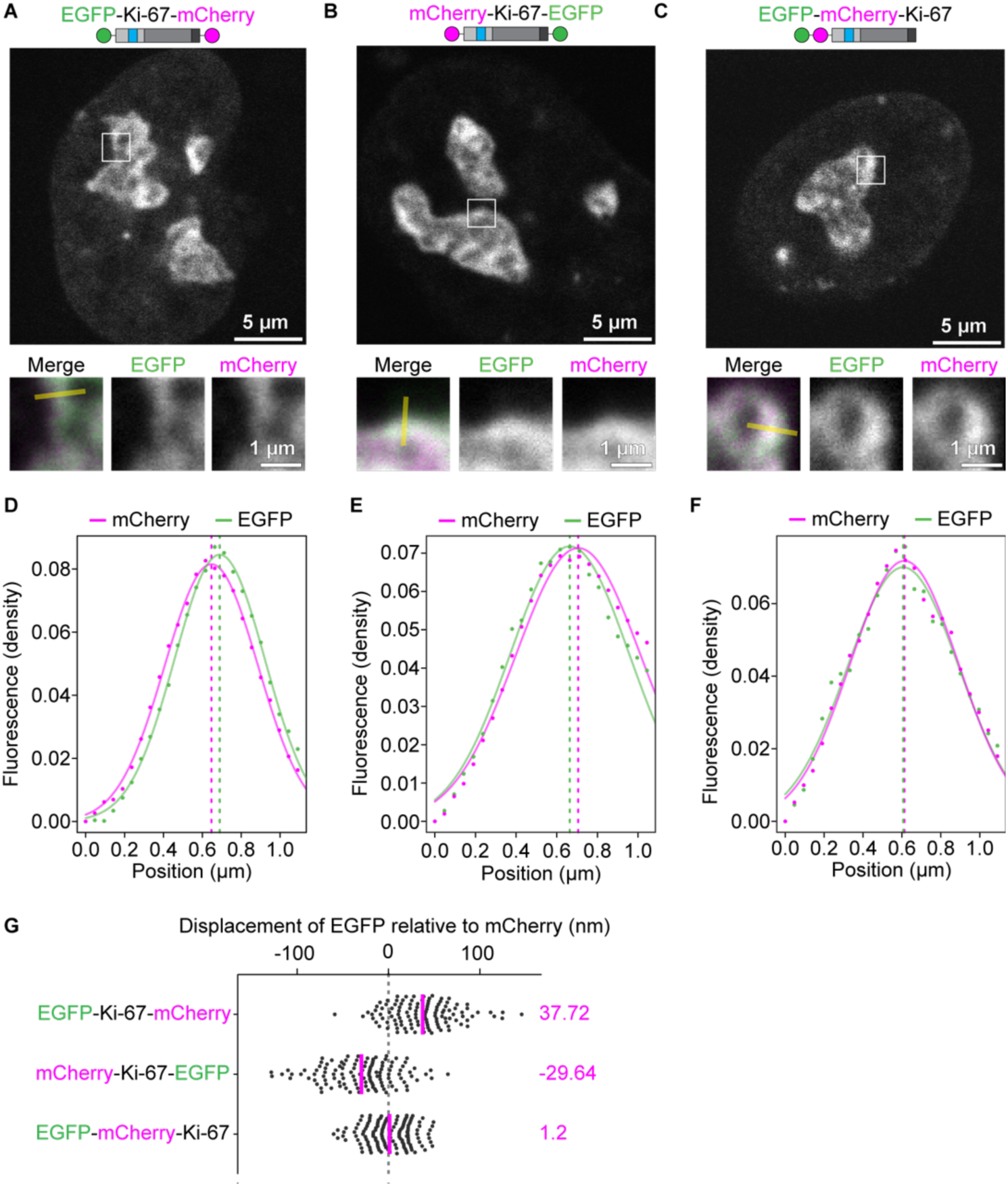
Ki-67 adopts a partially extended, preferentially oriented structure within the nucleolus. (A–C) Live cell imaging of dual fluorescence-tagged Ki-67. Cells were transfected with EGFP-Ki-67-mCherry (A), mCherry-Ki-67-EGFP (B), or EGFP-mCherry-Ki-67 (C) and imaged using a confocal microscope. Insets display EGFP and mCherry signals at the nucleolar rim. The yellow line indicates the line profile from outside to inside the nucleolus. (D–F) Quantification of fluorescence signal densities along the line profile. Fluorescence intensity values were extracted along the indicated line profiles and fitted with Gaussian functions (solid lines). The positions of peak fluorescence intensities derived from the Gaussian fits are marked with dashed lines. (G) Quantification of EGFP signal displacement relative to mCherry signals at the nucleolar periphery. Line profiles were performed in cells transiently expressing EGFP-Ki-67-mCherry, a reversed order of fluorescent proteins (mCherry-Ki-67-EGFP), or N-terminal tagging of both fluorescent tags (EGFP-mCherry-Ki-67). Displacement was measured as the distance between EGFP and mCherry peaks after Gaussian fitting. For (G), n = 116 lines from 52 nuclei (EGFP-Ki-67-mCherry), n = 119 lines from 52 nuclei (mCherry-Ki-67-EGFP), n = 135 lines from 53 nuclei (EGFP-mCherry-Ki-67), 3 experiments.

In summary, we have shown that Ki-67 localises specifically to the chromatin-nucleolar boundary with a clear orientation and an extension of ∼33 nm. The dual affinity domains, CP and LR, separated by a repeat-containing region, are essential for Ki-67 boundary localisation and regulation of nucleolar shape and nucleolar chromatin enrichment.

### Ki-67-dependent chromatin recruitment regulates nucleolar shape

Given that our previous analysis showed a strong link between irregular nucleolar shape and increased chromatin incorporation within nucleoli, we hypothesised that the chromatin network within and around nucleoli might be the major determinant of nucleolar shape.

To test this, we revisited our initial screening hits and aimed to test whether nucleolar rounding induced by knockdown of the top 20 candidate genes correlates with chromatin depletion both within nucleoli and at the rim. We depleted candidate genes with two to three different siRNAs in cells stably expressing H2B-mCherry and SNAP-NPM1 and imaged cells using automated confocal spinning disk microscopy. Strikingly, the depletion of all 20 candidate proteins led to a reduction in chromatin levels within the nucleolar interior resembling the effects of Ki-67 depletion (Figure 7A and S8A). Additionally, the decrease in chromatin levels within the nucleolar rim was evident in the majority of these conditions (Figure 7B). This finding supports the hypothesis that round nucleoli are generally depleted of chromatin.

**Figure 7.**
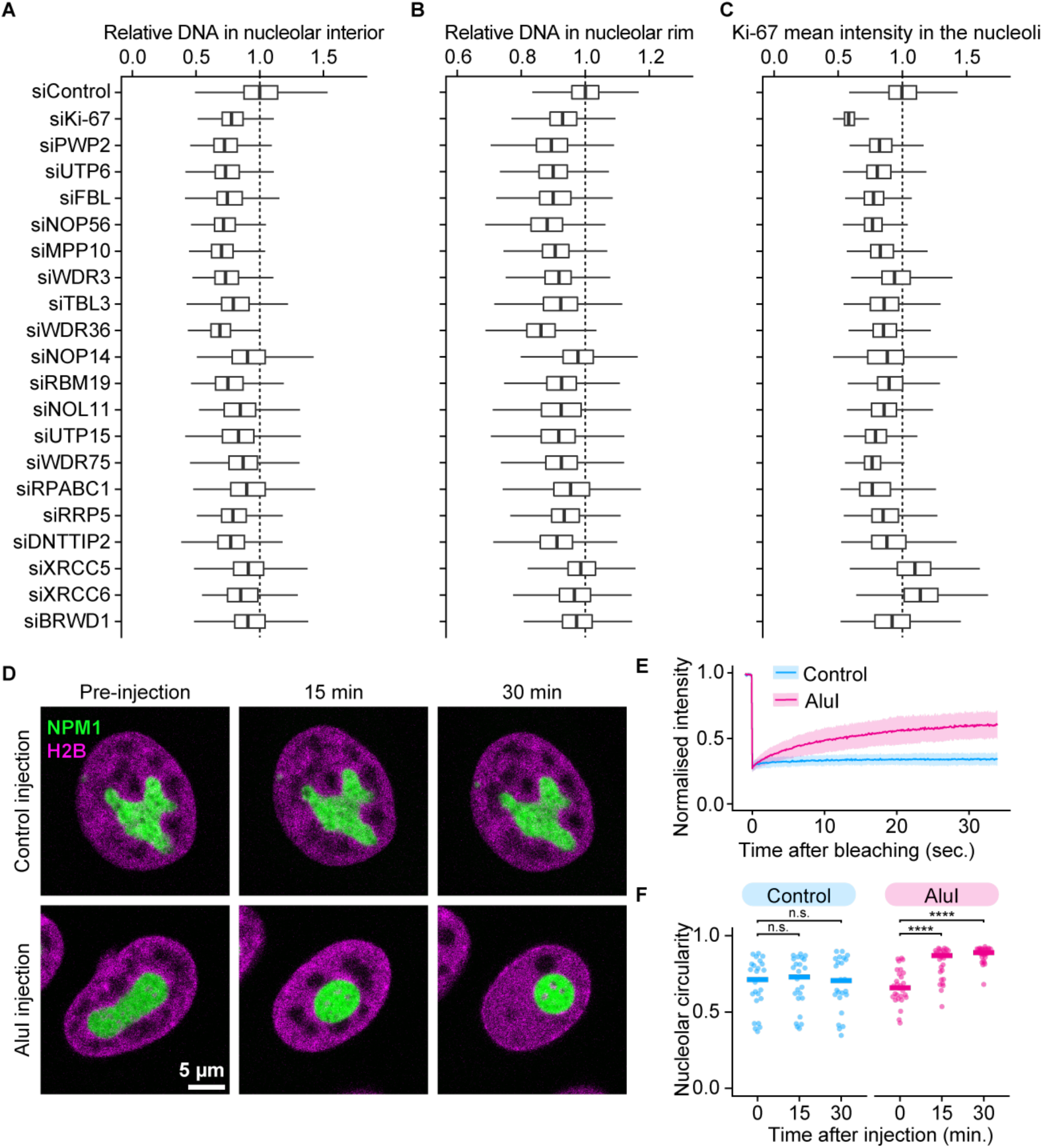
The chromatin network in and around nucleoli is a key determinant of nucleolar shape. (A, B) Quantification of chromatin levels in the nucleolus following the depletion of the top 20 candidates causing nucleolar rounding (Figure 1). Images of HeLa cells stably expressing NPM1-EGFP and H2B-mCherry were acquired using spinning disk microscopy. Relative H2B signal intensities in the nucleolar interior and the nucleolar rim were calculated as described in (Figure 3D and 3E). (C) Quantification of nucleolar Ki-67 levels following the depletion of the top 20 candidates causing nucleolar rounding (Figure 1). Images of endogenous EGFP-Ki-67 HeLa cells stably expressing SNAP-NPM1 were acquired using a wide-field microscope. The mean intensity of Ki-67 signals was measured in nucleoli, and median values per nucleus were calculated. (D) Effect of AluI injection on the nucleolus. HeLa cells stably expressing SNAP-NPM1 and H2B-mNeongreen were injected with a control buffer or an AluI-containing buffer. (E) Fluorescence recovery curves of H2B-mNeonGreen following AluI injection were compared to those of undigested cells. Lines and shades indicate mean ± SD. (F) Quantification of the nucleolar shape 30 min after AluI injection. The median nucleolar circularity per nucleus is shown. Bars indicate the median. Lines indicate mean values. For (A), n > 1000 cells per candidate. 3 experiments. For (B–D), n > 500 cells per candidate. 3 experiments. For (E), n = 15 nucleus (undigested) and n = 14 (AluI injection), 2 experiments. For (F), n = 25 nuclei (Control injection) and n = 29 (AluI injection), 2 experiments. **** P < 0.0001 with Wilcoxon Test. See also Figure S8.

To investigate whether Ki-67 plays a crucial role in chromatin depletion under these conditions, we knocked down candidate genes in a cell line stably expressing endogenous EGFP-Ki-67 and stable overexpressing SNAP-NPM1, followed by imaging with automated widefield microscopy. Interestingly, in 18 of the 20 knockdowns, Ki-67 levels were lower than in the control (Figure 7C and S8B), raising the possibility that Ki-67 may regulate the chromatin environment even in these conditions. However, nucleolar rounding and chromatin depletion from nucleoli can occur even when Ki-67 levels remain high. For instance, depletion of the DNA damage proteins XRCC5 and XRCC6 increased Ki-67 levels but still led to rounded nucleoli and a marked reduction of intra-nucleolar chromatin. In summary, these findings suggest that chromatin is the major determinant of nucleolar shape.

As an alternative test of this conclusion, we aimed to acutely disrupt the chromatin network within and around nucleoli and observe the effect on nucleolar shape. We performed acute chromatin digestion in a cell line expressing a nucleolar marker (SNAP-NPM1) and a chromatin marker (H2B-mNeonGreen) by microinjecting the restriction enzyme AluI (Figure 7D), which has been shown to efficiently digest chromatin in living cells^51^. We confirmed successful chromatin digestion by observing an increase in H2B-mNeonGreen mobility (Figure 7E). Remarkably, within 15 min of AluI microinjection, nucleoli became noticeably rounded (Figure 7D and 7F). Together, these findings reinforce the hypothesis that the chromatin network plays a central role in determining nucleolar shape.

## Discussion

Here, we uncover a chromatin-based mechanism that regulates nucleolar shape (Figure 8): Ki-67 recruits chromatin to both the surface and the interior of nucleoli, causing them to adopt an irregular shape. This chromatin tethering function relies on Ki-67’s dual affinity for chromatin and nucleoli: The N-terminal CP domain has affinity for nucleoli, while the C-terminal LR domain, separated from the CP by a repeat region, interacts with chromatin. This dual interaction drives Ki-67 localisation at the chromatin-nucleolus interface in a preferred molecular orientation, promoting chromatin enrichment both inside and around nucleoli. Given that chromatin depletion from the nucleolus is commonly observed in rounded nucleoli (Figure 7A and 7B), we propose that the chromatin environment of nucleoli plays a crucial role in regulating nucleolar shape. When chromatin is tightly tethered to the nucleolus, it affects the rigidity and flexibility due to the viscoelastic properties of chromatin, which may deform or stabilise the shape of nucleoli.

**Figure 8.**
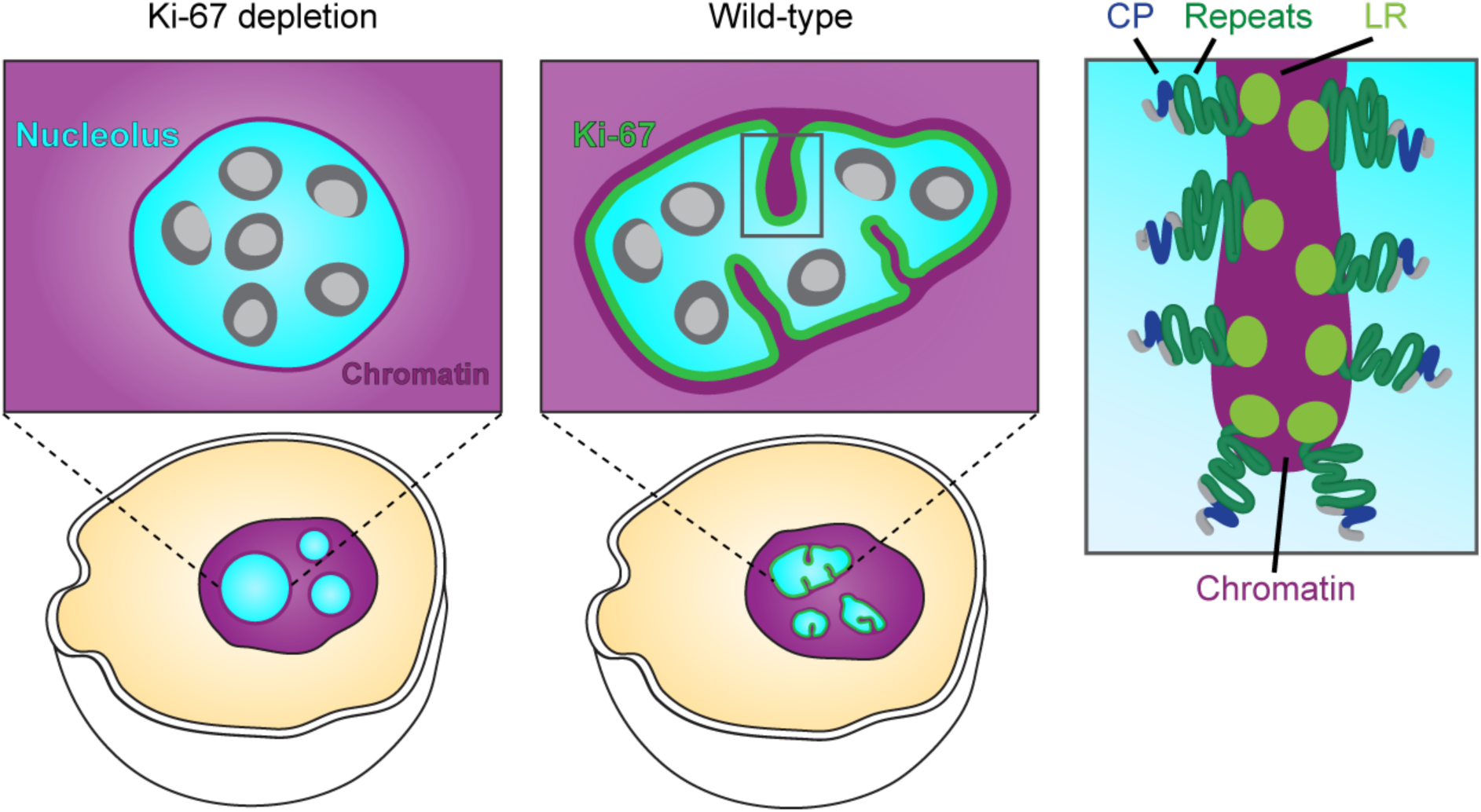
Ki-67-mediated chromatin anchoring to the nucleolus shapes nucleolar morphology. Working model of nucleolar shape regulation by Ki-67. Ki-67 anchors chromatin in the nucleolar interior and the rim due to its amphiphilic nature, characterised by a positively charged patch (CP) that interacts with the nucleolus and a leucine-arginine-rich (LR) domain that binds to chromatin, separated by a repeat domain. This chromatin network within and around the nucleolus prevents nucleolar rounding and thus determines nucleolar shape.

Previous data revealed that Ki-67 functions as a biological surfactant^35^, separating mitotic chromosomes by localising to their surface and adopting an extended structure (∼90 nm), mediated by Ki-67’s amphiphilic nature towards both chromatin and cytoplasm. Given its ability to promote non-spherical nucleolar shapes and solubilises chromatin within nucleoli, it is tempting to speculate that Ki-67 may also behave as a surfactant-like protein during interphase. However, our previous findings indicate that Ki-67 loses its surfactant-like behaviour during mitotic exit: Dephosphorylation triggers both Ki-67’s phase separation with RNA and its collapse to a ∼30 nm structure, promoting chromosome adhesion instead of repulsion^36,37^.

Here, we show that Ki-67 remains in its collapsed form during interphase (Figure 6) and, interestingly, displays patchy localisation at the chromatin-nucleolus interface (Figure 2A), suggesting it remains in a phase separated state. Additionally, its interphase mobility, as measured by fluorescence recovery after photobleaching (FRAP), is markedly reduced compared to mitotic Ki-67^47^, suggesting that its interactions are more stable and not characteristic of dynamic, surfactant-like behaviour^52^. Together, these observations suggest that Ki-67 may operate through a distinct mechanism during interphase compared to it surfactant-like role on mitotic chromosomes^35^.

Instead, Ki-67 appears to behave more like a Pickering agent composed of solid-like particles that adsorb at interfaces and stabilise emulsions^53^. Such agents have been described in P-granules, membrane-less organelles in *C. elegans*^54,55^, where MEG-3 protein clusters exhibit slow dynamics and solid-like characteristics. While direct measurements of nucleolar surface tension and the fusion dynamics of intra-nucleolar chromatin foci are lacking, Ki-67’s reduced molecular extension, patchy localisation and low mobility during interphase are consistent with Pickering agent-like behaviour.

As an interfacial stabiliser, Ki-67 not only shapes nucleolar morphology but also influences the organisation of surrounding chromatin, which is closely associated with heterochromatin^8^. Supporting this view, previous studies have shown that Ki-67 contributes to the maintenance of heterochromatin. During interphase, Ki-67 depletion leads to chromatin decompaction, particularly in perinucleolar regions, and a significant decrease in H3K9me3 levels within heterochromatic regions^34^. Conversely, overexpression of Ki-67 caused ectopic heterochromatin formation^34^. Consistent with these observations, Ki-67 preferentially localises to late-replicating genomic regions at the nuclear periphery and around the nucleolus, which are enriched in H3K9me3^42^. Moreover, Ki-67 is required for tethering heterochromatic elements to the nucleolus, such as satellite DNA^56^, a LacO array near rRNA repeats on chromosome 13^49^, centromeric regions marked by CENP-A^34^, and the inactive X chromosome^57^. Overall, loss of Ki-67 disrupts the positioning of these heterochromatic regions, leading to their relocalisation.

Despite the clear role of Ki-67 in heterochromatin formation and spatial positioning of genomic regions, its impact on gene expression remains less well defined. While several studies reported wide-spread gene deregulation following Ki-67 depletion^34,57,58^, others observed minimal changes in transcription, with gene repression largely unaffected in HCT116 cells^42,59^. This discrepancy may reflect differences in cell type, particularly whether the p21 pathway is active, as p21 induction following Ki-67 loss can cause secondary effects on gene expression due to cell-cycle perturbations^57^.

In addition, lamina-associated and nucleolus-associated heterochromatin can dynamically exchange^10,60,61^, suggesting that loss of nucleolar tethering may redirect chromatin to the nuclear periphery, where it can remain transcriptionally silent. Consistent with this, upon Ki-67 depletion, regions previously bound by Ki-67 show increased Lamin interactions^42^. This spatial redistribution may help preserve the transcriptionally repressive state of these regions, which could explain the relatively modest changes in gene expression observed upon Ki-67 loss. Thus, Ki-67 appears to be one of several overlapping mechanisms that preserve heterochromatin by regulating its spatial positioning.

The precise mechanism by which Ki-67 regulates heterochromatin remains incompletely understood. Its effects may be mediated through previously described direct interactions with chromatin-associated proteins such as HP1^62,63^ or the CAF-1 complex^56,64^. Our data, including the observed amphiphilic nature of Ki-67 and the excessive chromatin incorporation into nucleoli upon overexpression, support a model in which Ki-67 primarily serves a structural role. We propose that Ki-67 acts as a scaffold that tethers chromatin to the repressive environment of the nucleolus, thereby promoting compaction and silencing indirectly through spatial sequestration. Whether and how Ki-67 contributes more directly to the formation or maintenance of heterochromatic marks remains to be fully explored.

By anchoring chromatin to the nucleolar surface and interior, Ki-67 promotes the formation of irregularly shaped nucleoli with increased surface area. This expanded interface may not only facilitate transcriptional repression, but may also enhance the directional export of mature ribosomal subunits to the cytoplasm, supporting the elevated ribosome production necessary for cell growth and protein synthesis. Intriguingly, rapidly proliferating cells, such as stem/progenitor cells and cancer cells, generally express high levels of Ki-67^65,66^ and are characterised by large, irregular nucleoli^18,19^, features associated with their high proliferative and metabolic demands^67^. Conversely, Ki-67 protein is typically undetectable in senescent cells^68,69^. A hallmark of cellular senescence is a change in cell morphology^70^, which includes the nucleolus adopting a more rounded shape^71^ along with loss of H3K9me3 and displacement of centromeric and pericentromeric alpha-satellite sequences^72^ – regions that strongly interact with Ki-67^42^. These changes align with our findings that loss of Ki-67 causes nucleolar rounding and disrupts chromatin tethering to nucleoli. These observations suggest that nucleolar shape may be functionally linked to both chromatin organisation and cellular growth states, including their impact on ribosome production and gene expression.

Our study further highlights the importance of interfacial proteins in tightly connecting and shaping two membrane-less compartments, nucleoli and heterochromatin. In light of the growing recognition of the interfaces between biological condensates and cellular structures, including chromatin, RNA, and other membrane-less compartments^73,75,77,79,81^, amphiphilic proteins like Ki-67 might be crucial in controlling the properties and fluxes^77–80^ between membrane-less compartments, thereby influencing their organisation and function. Further studies of interfacial assemblies in cells, as well as on synthetic co-condensates in vitro^82^, will not only enhance our understanding of the regulatory mechanisms that control phase separation in cells but will also inform the rational design of engineered condensates, with potential applications in biosynthesis and drug delivery.

## Acknowledgements

We thank Daniel W. Gerlich for providing cell lines, the EMBL Advanced Light Microscopy Facility (ALMF) for imaging support, and Christian Tischer and Jean-Karim Hériché for assistance with the custom image analysis. This work was supported by the German Research Foundation (DFG project number 402723784) and the Human Frontier Science Program (CDA00045/2019). A.H.-A. has received a PhD fellowship from the Boehringer Ingelheim Fonds; Y. H. was supported by a fellowship from the EMBL interdisciplinary Postdoc (EIPOD) program (Marie Sklodowska-Curie Actions, COFUND grant agreement 664726).

## Author contributions

D.S., Y.H., and S.C.-H. conceived the project. D.S., Y.H., L.F. and M.C. designed, performed and analysed all experiments. A.H.-A. performed initial experiments on Ki-67 mutants’ localisation. S.C.-H. acquired funding. B. N. supported an experimental set-up for a high throughput screen. S.C.-H. supervised the project. D.S., Y.H., A.H.-A, and S.C.-H. wrote the manuscript.

## Declaration of generative AI and AI-assisted technologies in the writing process

During the preparation of this work, the authors used ChatGPT and Claude in order to enhance clarity, improve readability and eliminate redundancy. After using this tool/service, the authors reviewed and edited the content as needed and take full responsibility for the content of the publication.

## Materials and Methods

### Cell lines and cell culture

All of the cell lines used in this study have been regularly verified as negative for mycoplasma contamination. Their sources and authentication are summarised in Supplementary Table 2. All cell lines used in this study were HeLa-Kyoto cells that have been previously described^83^ . Cells were cultured in Dulbecco’s modified medium (DMEM; Thermo Fisher Scientific, 41965039) containing 10% (v/v) fetal bovine serum (FBS; Thermo Fisher Scientific, 10270106), 1% (v/v) penicillin-streptomycin (Sigma Aldrich, 15140122), 1 mM Sodium Pyruvate (Thermo Fisher Scientific, 11360039). For drug selection, cells were cultured in the above medium supplemented with antibiotics according to the expression constructs: geneticin G418 sulphate (Thermo Fisher Scientific, 11811031) at a final concentration of 300 µg/mL, puromycin (Merck Millipore, 540411) at a final concentration of 0.5 µg/mL, blasticidin S hydrochloride (Sigma Aldrich, 15205) at a final concentration of 6 µg/mL, and hygromycin S at a final concentration of 300 µg/mL (Thermo Fisher Scientific, 10687010). For live-cell imaging, cells were cultured in FluoroBrite DMEM (imaging medium; Thermo Fisher Scientific, A1896701) containing 10% (v/v) FBS, 1% (v/v) penicillin-streptomycin, 1 mM sodium pyruvate, and 1% (v/v) GlutaMAX (Thermo Fisher Scientific, 35050038). All live imaging experiments were performed under the condition of constant humidity and 37°C temperature with 5% CO_2_.

### Plasmid construction

All constructs are listed in Supplementary Table 3. For lentivirus transfer vectors, protein-coding sequences and protein tags were inserted into a backbone vector containing internal ribosome entry site (IRES) and antibiotics genes by Gibson assembly. To generate the donor vector for EGFP-miniDegron-Ki-67 knock-in, the miniDegron sequence^45^ was inserted into an EGFP knock-in donor vector for Ki-67 N-terminus^35^ by Gibson assembly. For the donor vector for the Halo-tag knock-in at the UBF N-terminus, homology arm sequences (700 bp around the start codon of UBTF gene) and the Halo-tag sequence were synthesised and cloned into pUC57 vector (Biomatik). To generate single-guide RNAs (sgRNAs)/Cas9 nickase (Cas9 D10A) expressing plasmids, sgRNAs were designed on Benchling (https://www.benchling.com/) and assembled into px335 vector (Addgene #42335). The following sgRNA sequences were used: sgRNA Ki-67 no. 1: 5’-AATGTGGCCCACGAGAC-GCC-3’, sgRNA Ki-67 no. 2: 5’-tgagtataatccgtagggga-3’, sgRNA UBTF no. 1: 5’-AGCCAC-CTCCTCGGTCGTGC-3’, sgRNA UBTF no. 2: 5’-AGCCGACTGCCCCACAGACC-3.

### Generation of stable cell lines

Cell lines stably expressing for fluorescence labelled marker proteins were generated by random plasmid integration or a lentiviral vector system pseudotyped with a VSV-G or a mouse ecotropic envelope that is rodent-restricted (RIEP receptor system). Construction of RIEP receptor parental cell lines and subsequent generation of stable cell lines that express fluorescent marker proteins were performed as described previously^84^. Genome editing was performed by the CRISPR-Cas9 nickase approach as described^85^. HeLa cells endogenously tagged with EGFP at the N-terminus of Ki-67 were previously described^35^. For the generation of auxin-inducible degradation of Ki-67, two sgRNAs targeting Ki-67 and the donor vector containing miniDegron-EGFP having 1-kb homology arms were transfected into cells using X-tremeGENE 9 DNA transfection reagent (Roche, 6365779001). Following 1-week culture, GFP-positive cells were sorted into a 96-well plate using FACSAria™ Fusion Flow Cytometer (BD Bioscience). Expanded cells were assessed by light microscopy for cell morphology, followed by genotyping, immunoblotting, and GFP signal localisation by confocal fluorescence microscopy. Following the validation of homozygous knock-in cells, OsTiR1 and FBL-TagRFP were stably expressed in the miniDegron-EGFP knock-in cells by lentivirus transduction. For the generation of endogenously expressing Halo-UBF, cells were electroporated with the two Cas9 nickase–sgRNA-expressing plasmids, 5 μg each, and 7.5 μg Halo-tag knock-in donor plasmid using the Neon Transfection System (Thermo Fisher Scientific) with 3 × 10 ms pulses at 1,300 V. Following 1-week culture, cells were incubated with 50 nM Halo-TMR for 30 min before sorting. Cells showing TMR positive signals were sorted into a 96-well plate using FACSAria™ Fusion Flow Cytometer (BD Bioscience). Expanded cells were assessed by light microscopy for cell morphology, followed by genotyping, immunoblotting, and Halo-UBF signal localisation via fluorescence microscopy.

### RNAi screen

For the screen on nucleolar shape (Figure 1), 614 genes were targeted by either two or three Silencer select siRNAs (Thermo Fisher Scientific). This target gene list included 63 nucleolar proteins localising to the chromosome surface during mitosis^86^, 139 genes involved in regulating nucleolar number^22^, 112 proteins whose solubility changes in response to increased ATP levels^24^, and 310 proteins that undergo solubility changes during mitosis^23^ . Some genes were present in more than one library. Using solid-phase reverse transfection^87^, 384-well imaging plates (Cellvis, P384-1.5H-N) were coated with siRNA transfection mixes. Cells stably expressing NPM1-EGFP and H2B-mCherry were seeded on the screening plates 72 h prior to imaging using a Multidrop Reagent Dispenser (Thermo Scientific). Images were acquired at 4 different positions in each well on an ImageXpressMicro XL screening microscope (Molecular Devices) using a ×20, 0.75 NA CFI P-Apo Lambda objective (Nikon).

For the follow-up experiments on nucleolus-associated chromatin levels and Ki-67 levels (Figure 7A–7C), siRNAs transfection and cell seeding were prepared with the same approaches using a subset of target genes in 96-well imaging plates (Cellvis, P96-1.5H-N). For the nucleolar chromatin analysis (Figure 7A, 7B, and S8A), Cells stably expressing NPM1-EGFP and H2B-mCherry were acquired at 4 different positions in each well on an IXplore SpinSR spinning disc microscope (Olympus) with a CSU-W1-TS 2D 50/SoRa confocal scanner (Yokogawa) using UPLXAPO 20X objective (Olympus). For the Ki-67 levels analysis (Figure 7C and S8B), cells expressing endogenously tagged EGFP-Ki-67 and stably expressing SNAP-NPM1 and H2B-mCherry were seeded to 384-well imaging plates. SNAP-NPM1 were labelled with 25 nM SNAP-SiR (New England Biolabs, S9102S). Images were acquired at 4 different positions in each well on an ImageXpressMicro XL screening microscope using a ×20, 0.75 NA CFI P-Apo Lambda objective.

### Protein knock-down by siRNA transfection

Pre-designed SilencerSelect siRNAs (Thermo Fisher Scientific) were diluted in nuclease-free water (Thermo Fisher Scientific) to a concentration of 10 µM as a stock solution. For fluorescence imaging, the stock siRNA solution was further diluted to 80 nM in DEPC-treated water (Thermo Fisher Scientific) for a working solution. In an 8-well Lab-Tek chamber slide (Thermo Fisher Scientific, 155411), 50 µl of siRNA working solution was mixed with 50 µl of Opti-MEM containing 0.6 µl of Lipofectamine RNAiMAX Reagent (Thermo Fisher Scientific, 13778030). The cell suspension was incubated for 20 min at RT and seeded to achieve 50-70% confluency on the day of imaging. To assess the nucleolar roundness and the chromatin enrichment in the nucleolus, cells were imaged 72 h after siRNA transfection on a Zeiss LSM780 using a Plan-Apochromat ×63/1.4 NA Oil DIC M27 oil immersion objective. To examine the effect of Ki-67 depletion on the nucleolar small subcompartment, cells were imaged 48 h after siRNA transfection on an Olympus iXplore SPIN SR using a UPLSAPO

×100/1.35 NA Silicon oil immersion objective. For holotomographic imaging, 2.5 µL of 10 µM siRNA stocks were diluted in 125 µL Opti-MEM and added to ibidi 35 mm high dishes (ibidi, 81156). Pre-mixed 7.5 µl of 125 µL of a transfection mix prepared from 125 µL per dish Opti-MEM and 7.5 µL/dish Lipofectamine RNAiMAX in 125 µL Opti-MEM was mixed with the diluted siRNA in the dishes. Twenty minutes after incubation at RT, cells suspended in the imaging medium were seeded to achieve 50-70% confluency on the day of imaging. Cells were imaged 72 h after siRNA transfection on 3D Cell Explorer microscope (Nanolive) at 37 °C with 5 % CO_2_ environment. The following siRNAs were used in this study; siControl (XWNeg9; s44426), sense strand 5’-UACGACCGGUCUAUCGUAGtt-3’, antisense strand 5’-CUACGAUAGACCGGUCGUAtt-3’; siKi-67 no. 1 (s8798), sense strand5‘-GUACCAUAAUAAUAGGGAAtt-3’; antisense strand UUCCCUAUUAUUAUGGUACaa siKi-67 no. 2 (s8796), sense strand 5’-CGUCGUGUCUCAA-GAUCUAtt-3’, antisense strand 5’-UAGAUCUUGAGACACGACGtg-3’

### Live cell Airyscan imaging

Cells were seeded on 8-well glass bottom slides (ibidi, 80807) one day before imaging. For Halo-tag and SNAP-tag labelling, cells were incubated with the medium containing 100 nM SNAP-SiR and 1×SPY595-DNA (Spirochrome, SC301) for 1 h at 37°C. After three washes with fresh medium, the cells were cultured in the imaging medium. Airyscan imaging was performed on a Zeiss LSM980 confocal microscope equipped with an Airyscan detector and a Plan-Apochromat ×63/1.4 NA Oil DIC M27 objective. Images were acquired using the Airyscan SR mode with 0.15 µm interval for 5 µm thickness. To correct for chromatic aberration, images of 100 nm TetraSpeck Fluorescent Microspheres (Thermo Fisher Scientific, T14792) were acquired using the same acquisition settings. Chromatic aberration was analysed from the bead images using Huygens profession. The mean chromatic aberration values of the beads were used to correct for chromatic aberration in the cell images.

### Immunofluorescence staining

Cells were seeded in a LabTek slide one day before fixation. To label Halo-tag and SNAP-tag, living cells were incubated with the medium 100 nM Halo-TMR and 100 nM SNAP-SiR (New England Biolabs, S9102S) for 20 min at 37°C. After three times washing with pre-warmed medium, cells were cultured in the fresh medium for 10 min. Then, cells were washed with phosphate-buffered saline (PBS) and fixed with 3.7% formaldehyde in PBS at room temperature (RT) for 15 min. After three times washing with PBS and quenching with 100 mM NH_4_Cl in PBS at RT for 10 min, cells were permeabilised with 0.2% (v/v) Triton X-100 in PBS at RT for 5 min. Followed by washing three times with PBS, blocking was performed with 3% Bovine serum albumin (BSA) in PBS at RT for 1 h, and then the cells were incubated with the primary antibody in the blocking buffer at 4°C overnight. The cells were washed three times with PBS and incubated with the secondary antibody in PBS at RT for 1 h. After twice washing with PBS, DNA was stained with 0.2 µg/ml DAPI (Thermo Fisher Scientific, 62248) in PBS at RT for 5 min. Images were acquired on a Zeiss LSM780 confocal microscope, operated by ZEN 2011, with an EC Plan-Neofluar ×40/1.30 Oil DIC M27 oil-immersion objective. The following antibodies were used: rabbit anti-Ki-67 antibody (Abcam, ab16667), goat anti-rabbit IgG conjugated with Alexa Fluor 488 (Thermo Fisher Scientific, A11034), goat anti-rabbit IgG conjugated with Alexa Fluor 594 (Thermo Fisher Scientific, A11037)

### Transient expression of dual fluorescence protein-tagged Ki-67

Wil-type cells were seeded to a LabTek slide and immediately transfected with plasmids coding EGFP-Ki-67-mCherry, mCherry-Ki-67-EGFP, and EGFP-mCherry-Ki-67 using polyethylenimine Max (PEI; Polysciences, 24765). Following two days of culture, images were acquired with a LSM780 confocal microscope and an EC Plan-Neofluar ×40/1.30 Oil DIC M27 oil-immersion objective operated by ZEN 2011. Single-cell z-stack images were acquired with EGFP channel and selected a region of the nucleolar periphery, preferentially at positions with minimal curvature in the z-dimension. Then, using zoom ×35, both EGFP and mCherry channels were acquired. Chromatic aberration was determined by acquiring images of TetraSpeck Fluorescent Microspheres with identical imaging settings as for nucleoli and measuring the average offset of intensity peaks in the x- and y-dimensions. The alignment of the red and green channels of images of nucleoli was then corrected for chromatic aberration accordingly.

### Ki-67 overexpression in cells endogenously expressing EGFP-Ki-67

A plasmid coding EGFP-Ki-67 was transfected in cells endogenously expressing EGFP-Ki-67 with PEI in a 6 cm dish and incubated for 1 day. Cells expressing higher levels of EGFP-Ki-67 compared to the endogenous EGFP-Ki-67 were sorted into 500 µL of the culture medium in a 1.5 mL tube and seeded in LabTek slides. Following 1-day culture, DNA was stained with 1 x SPY555-DNA (Spirochrome, SC201) in the imaging medium for at least 2 h. Imaging was performed on a Zeiss LSM780 microscope with an EC Plan-Neofluar ×40/1.30 Oil DIC M27 oil-immersion objective using custom-made macro in Zen Black for autofocus acquisition.

### Rescue the Ki-67 depletion phenotype with transient expression of Ki-67 mutants

Ki-67 mutant plasmids were transfected into Ki-67 KO cells in a 6 cm dish using PEI as described above. After culturing for 1 day, GFP-positive cells were isolated into 500 µl of the culture medium in a 1.5 mL tube using FACS. The sorting gate was determined by the EGFP levels of endogenously tagged EGFP-Ki-67 signal intensity. The sorted cells were spun down by 90× g for 3 min, resuspended in the imaging medium, and then seeded to LabTek slides. Following 1-day culture, DNA was stained with 0.2 µM SiR-DNA in the imaging medium for at least 2 h. Imaging was performed on a Zeiss LSM780 microscope with an EC Plan-Neofluar ×40/1.30 Oil DIC M27 oil-immersion objective using a custom-made macro in Zen Black for autofocus acquisition.

### Auxin-induced acute Ki-67 depletion and drug washout

Cells were seeded onto a 96-well imaging plate (Cellvis, P96-1.5H-N) within the imaging medium containing 1,250 cells. Following 1-day culture, DNA was stained with 0.2 µM SiR-DNA in the imaging medium 2 h before imaging. After equilibration of the imaging plate on the microscope stage at 37 °C for 1 h, time-lapse imaging was performed on a Zeiss LSM780 microscope with EC Plan-Neofluar ×40/1.30 Oil DIC M27 oil-immersion objective operated by Zen Black and a MyPic macro for autofocusing and acquiring multiple positions. Images were acquired with 3 z-slices by 0.525 µm intervals between slices at seven positions per well every 30 minutes. For the first time point, all wells remained untreated. After completion of the first time point acquisition for all wells at around 30 min after the beginning of time-lapse imaging, cells were treated with indole-3-acetic acid (IAA; abcam, ab146402) by replacing half the medium (100 µL out of total 200 µL per well) with imaging medium containing 500 µM IAA and 0.2 µg/ml SiR-DNA for the final concentration. For IAA non-treated cells, the 100 µL of the medium was replaced with 100 µL imaging medium only containing 0.2 µg/ml SiR-DNA for the final concentration. Around 8.5 hours after the beginning of time-lapse imaging, in one of the IAA-treated wells, IAA was washed out by replacing 150 µL of the medium with fresh imaging medium 10 times. Then, the 150 µL of the medium was replaced with 150 µL of imaging medium containing 0.2 µg/ml SiR-DNA. The time lapse was stopped after 22 h. For the IAA-insensitive cells, the time-lapse imaging was performed as described above.

### Microinjection

Live-cell microinjection experiments were performed using a FemtoJet microinjector (Eppendorf, 5181000017) in conjunction with an InjectMan NI2 micromanipulation device (Eppendorf, 5247000013). All microinjections were performed using pre-pulled Femtotips injection capillaries (Eppendorf, 5242952008). The microinjection device was directly mounted onto a customised confocal Zeiss LSM780, as described above.

Cells stably expressing H2B-mNeongreen and SNAP-NPM1 and endogenously Halo-tagged UBF were cultured in μ-Dish 35 mm high-wall imaging dishes with a polymer bottom (ibidi, 81156) to reach 80%-90% confluency on the day of the injection. SNAP-NPM1 and DNA were labelled with 100 nM SNAP-SiR and 0.2 μM SiR-DNA. AluI restriction enzyme (Thermo Fisher Scientific, FD0014) was mixed with twice the volume of microinjection buffer (50 mM HEPES pH 7.4 (home-made), 5% glycerol (Merck, 1.04092.2511), 1 mM Mg(OAc)_2_ (Sigma, M5661)) supplemented with 1 mg/mL CF®680R-conjugated 10k-MW dextran (VWR, 80116). Using an Eppendorf Microcapillary Microloader (Eppendorf, 5242956003), 4-6 μL diluted AluI in the microinjection buffer was loaded into a Femtotip. Microinjection was performed using 150 hPa injection pressure, 0.2 s injection time and 30 hPa compensation pressure. Images were acquired before injection, 15 min and 30 min after the injection on a Zeiss LSM780 using a Plan-Apochromat ×63/1.4 NA Oil DIC M27 oil immersion objective. For FRAP assay of H2B, photo-bleaching with 80% of 488 nm laser was performed 30 min after injection.

## Data analysis

### Image analysis of RNAi screen

Image analysis was performed in CellProfiler software^88,89^. For the nucleolar shape screen (Figure 1), the nuclei and the nucleoli were segmented by local adaptive thresholding. Using a control siRNA and siRNAs whose knock-down phenotypes are well known (siRNAs for INCENP, KIF11, PLK1, and CDC20), chromatin morphologies were analysed to assess the RNAi efficiency and specificity. These supervised datasets were used for an automatic classification method (Support Vector Machine with Radial Basis Function Kernel) to classify chromatin morphology in all images. Classification results were overlayed on images for quality control and identified mononucleated interphase cells with high confidence. The genes targeted by siRNAs show more than 20% of apoptotic cells were excluded from the following analysis. The median nucleolar aspect ratio was calculated in each well, followed by averaging those values for the same target genes except for control siRNA. For the control siRNA, the median value of the nucleolar aspect ratio in each control well was used.

For the analysis of the follow-up screen, we applied the same quality control methods and excluded cells other than interphase cells in the following analysis.

For the nucleolar chromatin analysis (Figure 7A, 7B, S8A), following segmentation of the nucleolus based on NPM1 signals, the segmentation was shrunk and expanded in arbitrary pixels to generate the nucleolar interior segmentation and the expanded nucleolus segmentation. The nucleolar rim segmentation was produced by subtracting the shrunk nucleolus segmentation from the expanded nucleolus segmentation. For the nucleoplasm segmentation, the expanded nucleolus segmentation was subtracted from the nucleus segmentation. To quantify the chromatin levels, H2B mean intensity was measured in each segmentation, followed by averaging those values per the nucleus for the same target gene.

### Segmentation of the nucleolus in fluorescence images

Single-slice images were selected based on the highest mean intensity of the nucleolar marker fluorescence signals across slices if the raw data contained a z-stack. The single-slice mages were analysed using CellProfiler ver. 3.1.8^88^ or ver. 4.2.5^89^. The nuclei and the nucleoli were segmented based on fluorescence signal intensity using an adaptive Otsu thresholding method. Based on the segmented nucleoli objects, the nucleolar interior segmentations were generated by shrinking arbitrary pixels from the nucleoli objects to exclude perinucleolar chromatin. The nucleolar rim segmentations were generated by expanding arbitrary pixels from nucleoli objects to cover the perinucleolar chromatin and then subtracting the original nucleoli masks. The nucleoli were linked to the parental nucleus.

Fluorescence signal intensity within the objects and the object size and shape were measured. The data were further analysed in R. Using the relationship between the parental nucleus and the children’s nucleoli, the average mean intensity of the nucleoli per nucleus was calculated. For the relative intensity measurements in the nucleolar interior and in the nucleolar rim, the average values of mean intensities in these segmentations were divided by the mean intensity in the parental nucleus. For the nucleolar area, the summed nucleolar area in the parental nucleus was calculated. For the nucleolar aspect ratio and circularity, the median value was calculated per the parental nucleus. The circularity was calculated as 4 * π * Area / Perimeter^2^.

To segment FC regions, UBF signals acquired by the spinning disk microscope were analysed using Pixel Classification in ilastik^90^. The classifier was manually trained on 10 images and then used for batch processing of the whole data sets to generate probability maps. Then, in CellProfiler, the FC regions was segmented based on the probability map with a 0.5 threshold. The segmented FC regions were linked to their parental nucleolus and nucleus, followed by the measurement of the median FC area and the FC density (the number of FC divided by the nucleolar area) per nucleolus in each nucleus.

### Radial signal distribution analysis

For the Airyscan imaging, the nucleolar segmentation was performed as previously described. To segment chromatin foci in the nucleolar interior, upper quantile values of the DNA intensity were measured within the nucleolar sub-region where the original nucleolus was eroded by 20 pixels. The upper quantile values were subtracted from the raw DNA signal intensity in the image. After applying the Gaussian filter on the image, chromatin foci were segmented by the adaptive Otsu method. The segmented chromatin foci were further filtered with their area and circularity between 3–15 pixels and above 0.7, respectively. Using a module called MeasureObjectIntensityDistribution, the expanded nucleolar mask was radially separated into 50 sectors from the chromatin foci in the nucleolar interior towards the edge of the expanded nucleolus and the mean intensity was measured within each sector. The sectors were divided into two regions: the chromatin foci (sector 1–15) and the nucleolar periphery (sector 16–50). In each region, min-max scaling was applied for each measured fluorescence signal in R.

For Ki-67 overexpression, the nuclei and the nucleoli were segmented as previously described. To measure radial signal distribution from the centre to the periphery of the nucleolus (Figure S5B and S5C), the expanded nucleolar mask was radially segmented into 10 sectors from the centre to the edge of the expanded nucleolar mask. In each fluorescence signal, min-max scaling was applied in R.

### Segmentation of nucleoli from holotomographic imaging data

Centre slices with in-focus nucleoli were manually chosen and isolated from raw 3-D holotomographic images. Background pixels and pixels belonging to nucleoli were manually annotated and trained in ilastik. Based on this training data, a neural network was developed by Akhmedkhan Shabanov^38^ and applied to segment nucleoli in the whole data sets. Nucleolar aspect ratio was measured in Fiji and analysed in R.

### Analysis of Ki-67 extension

To measure molecular extension, the fluorescence intensity of EGFP and mCherry signals was measured along line profiles of 2 pixels width drawn across the nucleolar periphery from the exterior to the interior across the nucleolar periphery in Fiji. The resulting intensity profiles were analysed in R. For each profile, background correction was performed by subtracting the minimum intensity value of each respective channel, EGFP and mCherry, from its raw data. These background-subtracted intensities for each channel were then normalized by dividing each value by the sum of all background-subtracted intensities within that channel for the profile. Subsequently, a single Gaussian function was fitted to each normalized channel profile to determine the Gaussian mean position. The displacement distance between EGFP and mCherry was then calculated as the difference between their respective Gaussian means.

### Colocalisation of Ki-67 and FBL or UBF

The nuclei and the nucleoli were segmented as described above. Pearson correlation coefficient of fluorescence signals of EGFP-Ki-67 and Halo-tagged UBF or FBL in the nucleoli was measured in CellProfiler. The data was further analysed in R.

### Analysis of Ki-67 mutant localisation

The nuclei and the nucleoli segmentation were performed as described above. The fluorescence intensity of Ki-67 mutants, NPM1 and DNA in the nucleolar rim and the nucleolar interior were plotted as scatter plots. Linear regression (*y* = *x* + 0) was applied to the data of full-length Ki-67tagged with EGFP. Using this linear regression model, the residuals were measured in each EGFP-tagged Ki-67 mutant, including the full-length Ki-67. The median and mean values of the residuals in each mutant were calculated and compared to the range of mean ± 2 SD of the full-length Ki-67 residuals. If both mean and median values in a mutant fall within this range, the mutant was considered to have similar localisation to the full-length Ki-67. The mutants with distinct localisation from the full-length Ki-67 were further analysed to assess their EGFP signal distribution within the nucleolus. The coefficient of variation (CV) of the mutant in the nucleolus was calculated by dividing the EGFP standard deviation intensity by the EGFP mean intensity.

### Statistical analysis

No statistical methods were used to predetermine the sample size. All experiments were repeated several times, and indicated experiment numbers always refer to biological replicates. Data were tested for normality and equal variances with Shapiro–Wilk and Levene’s tests (α= 0.05), respectively. To compare the two different groups, the appropriate statistical test was chosen as follows: Unpaired normal distributed data were tested with a two-tailed t-test (in case of similar variances) or with a two-tailed t-test with Welch’s correction (in case of different variances). Unpaired not normal distributed data were tested with a two-tailed Mann–Whitney test (in case of similar variances) or with a two-tailed Kolmogorov–Smirnov test (in case of different variances). For the multiple comparison, two-sided Dunn’s test was performed following Kruskal-Wallis test. All tests were performed with R or Prism.

---

**Figure S1.**
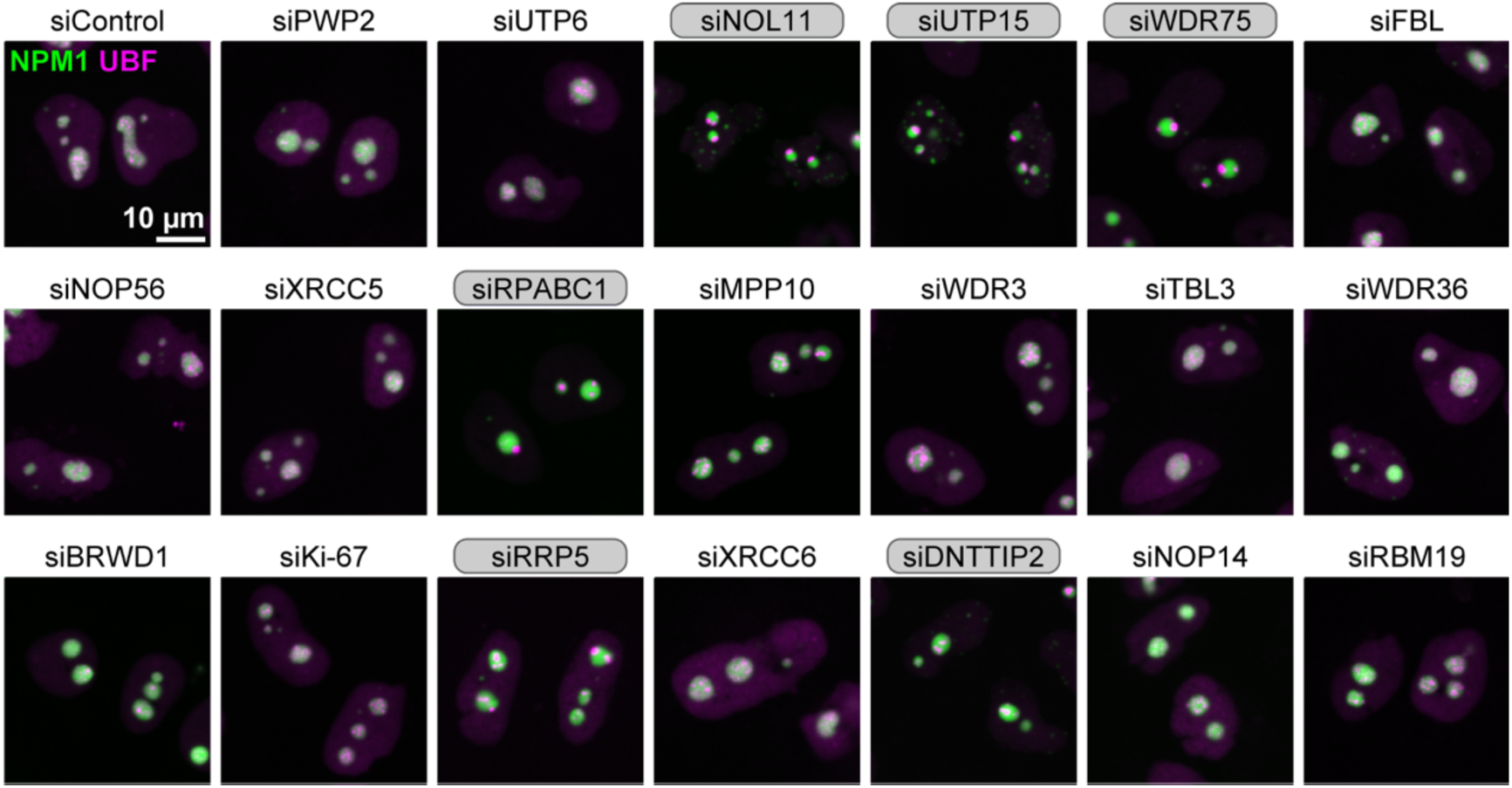
Depletion of several candidate genes that induce nucleolar rounding triggers nucleolar cap formation, related to Figure 1. Nucleolar cap formation following depletion of some top 20 candidates causes nucleolar rounding. HeLa cells endogenously tagged with Halo-UBF and stably expressing SNAP-NPM1 were transfected with a non-targeting control siRNA (siControl) and the siRNAs for the top 20 candidate genes (Figure 1). SNAP-NPM1 was labelled with SNAP-SiR (green), and Halo-UBF was labelled with Halo-Tetramethylrhodamine (TMR) (magenta). Nucleolar cap formation was manually annotated (grey boxes) based on the localisation of UBF.

**Figure S2.**
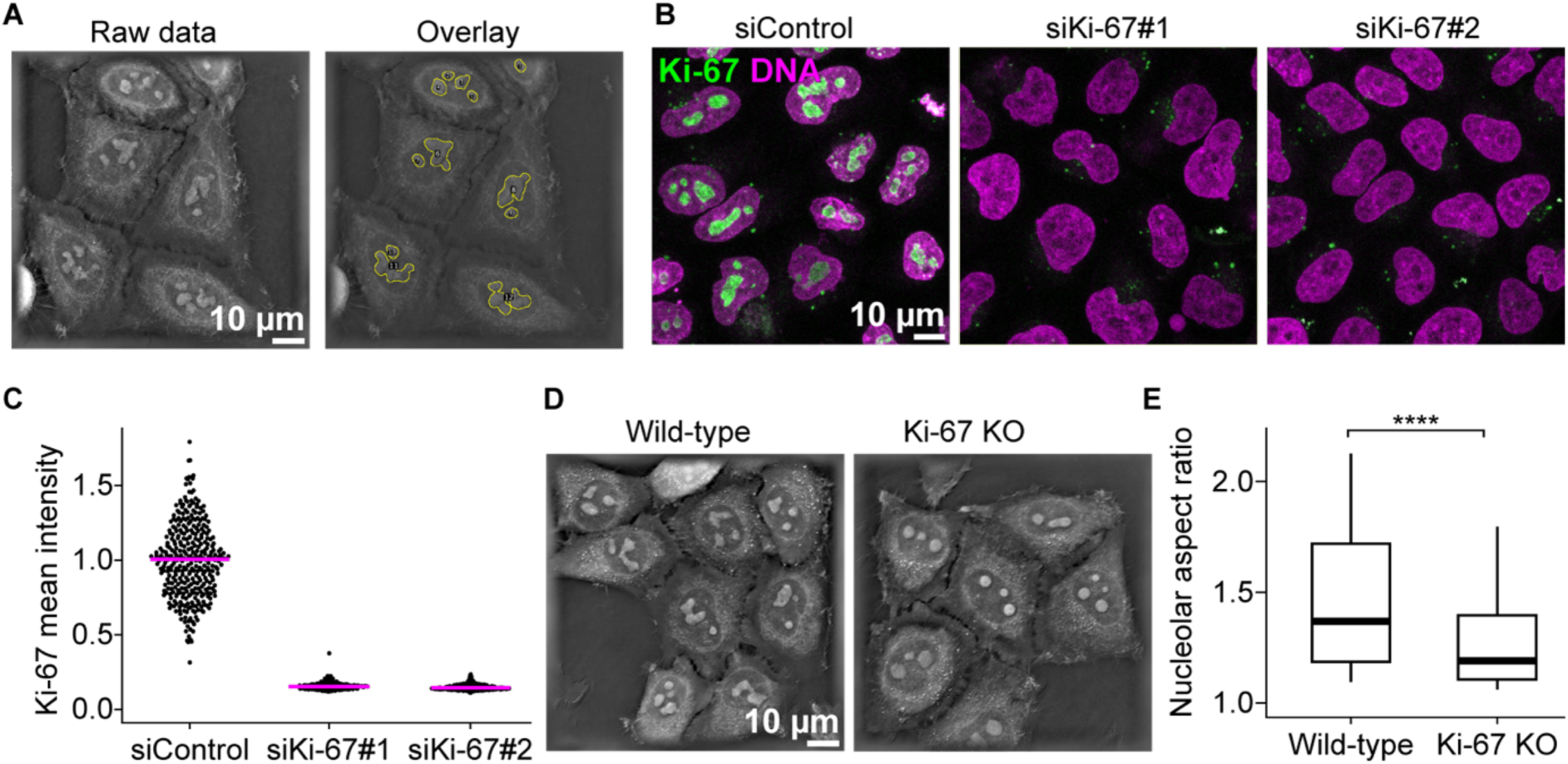
Ki-67 depletion or knockout leads to rounding, related to Figure 1. (A) Segmentation of nucleoli in holotomographic images. Raw holotomographic images (top) and overlay images of segmented nucleoli outlines (yellow) using a convolutional neural network on holotomographic images (bottom) are shown. (B, C) Confirmation of Ki-67 depletion by siRNAs. Cells expressing endogenously tagged EGFP-Ki-67 were transfected with two siRNAs for Ki-67. DNA was stained with SiR-DNA (B). EGFP-Ki-67 mean intensity in the nuclei was measured (B). (D) Label-free holotomographic live imaging of wild-type and Ki-67 knock-out (KO) cells. Single z-slices of a representative example are shown. (E) Quantification of nucleolar aspect ratio in holotomographic images. The aspect ratio of nucleoli was measured in wild-type and Ki-67 KO cells as in (F). Boxplots display the median (centre line), interquartile range (box), and whiskers extending to the 10th and 90th percentiles. For (E), n = 1407 nucleoli (wild-type cells), n = 1405 (Ki-67 KO), 3 experiments. ****P<0.0001 with Kolmogorov-Smirnov test.

**Figure S3.**
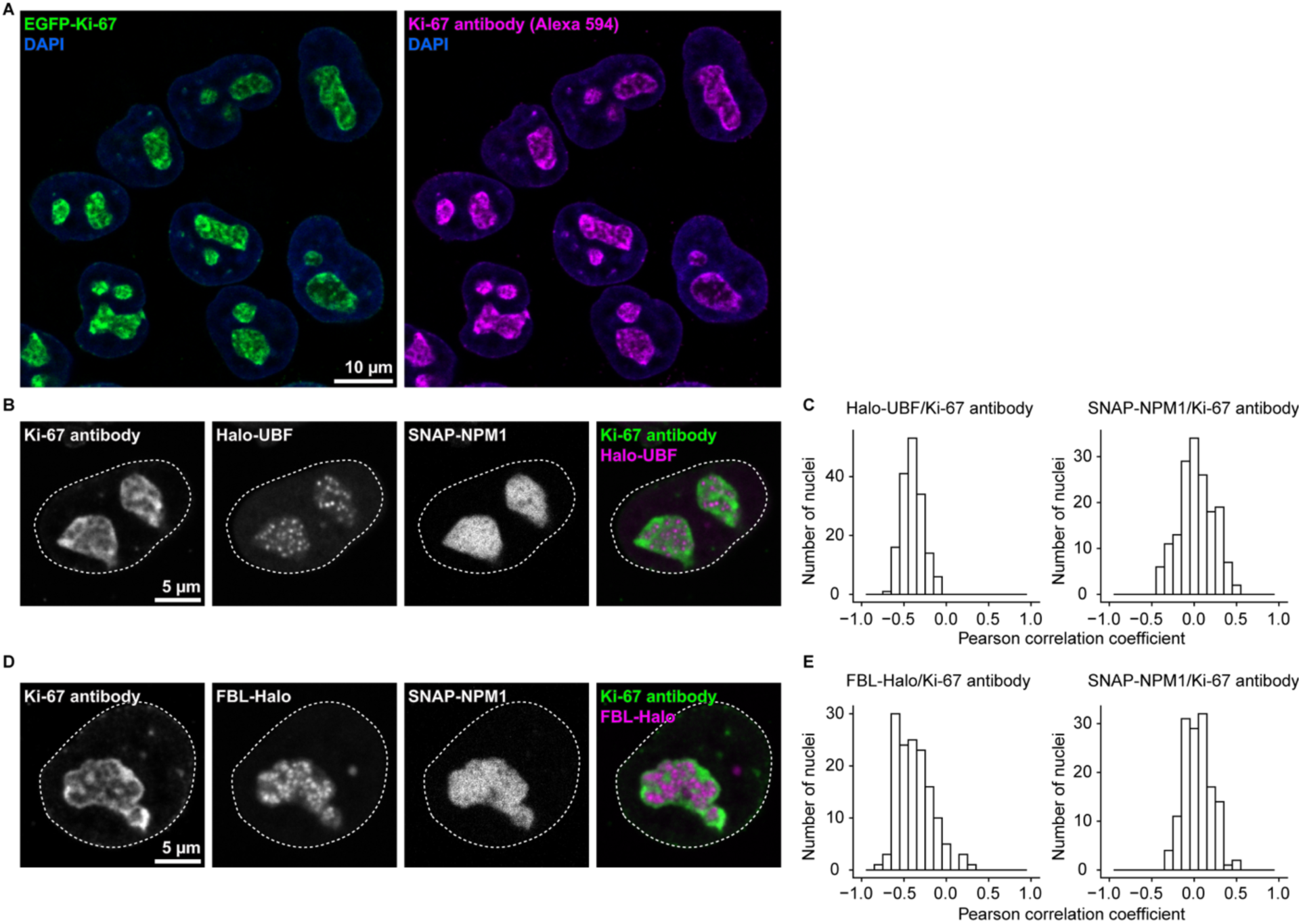
Ki-67 displays a distinct localisation pattern in the nucleolus, separate from the nucleolar subcompartments, related to Figure 2. (A) Immunofluorescence staining of Ki-67 in an endogenous EGFP-Ki-67 cell line. DNA was stained with DAPI. Notably, endogenous EGFP-Ki-67 signals substantially overlap with anti-Ki-67 antibody staining. (B) Co-staining of Ki-67 and UBF. After labelling cells expressing SNAP-NPM1 and Halo-UBF with SNAP-SiR and Halo-TMR in live cells, immunofluorescence was performed for Ki-67 as in (A). Dashed lines indicate the nuclear boundary. (C) Colocalisation analysis between Ki-67 and UBF or NPM1 within the nucleolus. Pearson correlation coefficient between Ki-67 and UBF signals (left panel) or Ki-67 and NPM1 signals (right panel) was measured within nucleoli (segmented based on NPM1 signal). A Pearson correlation coefficient of +1.0 indicates a perfect positive linear correlation, -1.0 indicates a perfect negative correlation, and 0 indicates no correlation between the signals. (D) Co-staining of Ki-67 and FBL. After labelling cells expressing SNAP-NPM1 and FBL-Halo with SNAP-SiR and Halo-TMR in live cells, immunofluorescence was performed for Ki-67 as in (A). Dashed lines indicate the nuclear boundary. (E) Colocalisation analysis between Ki-67 and FBL or NPM1 within the nucleolus. Pearson correlation coefficient between Ki-67 and FBL signals (left panel) or Ki-67 and NPM1 signals (right panel) was measured within nucleoli, segmented based on NPM1 signal as in (C). For (C–E), n = 144 nucleoli and n = 165 nucleoli, respectively.

**Figure S4.**
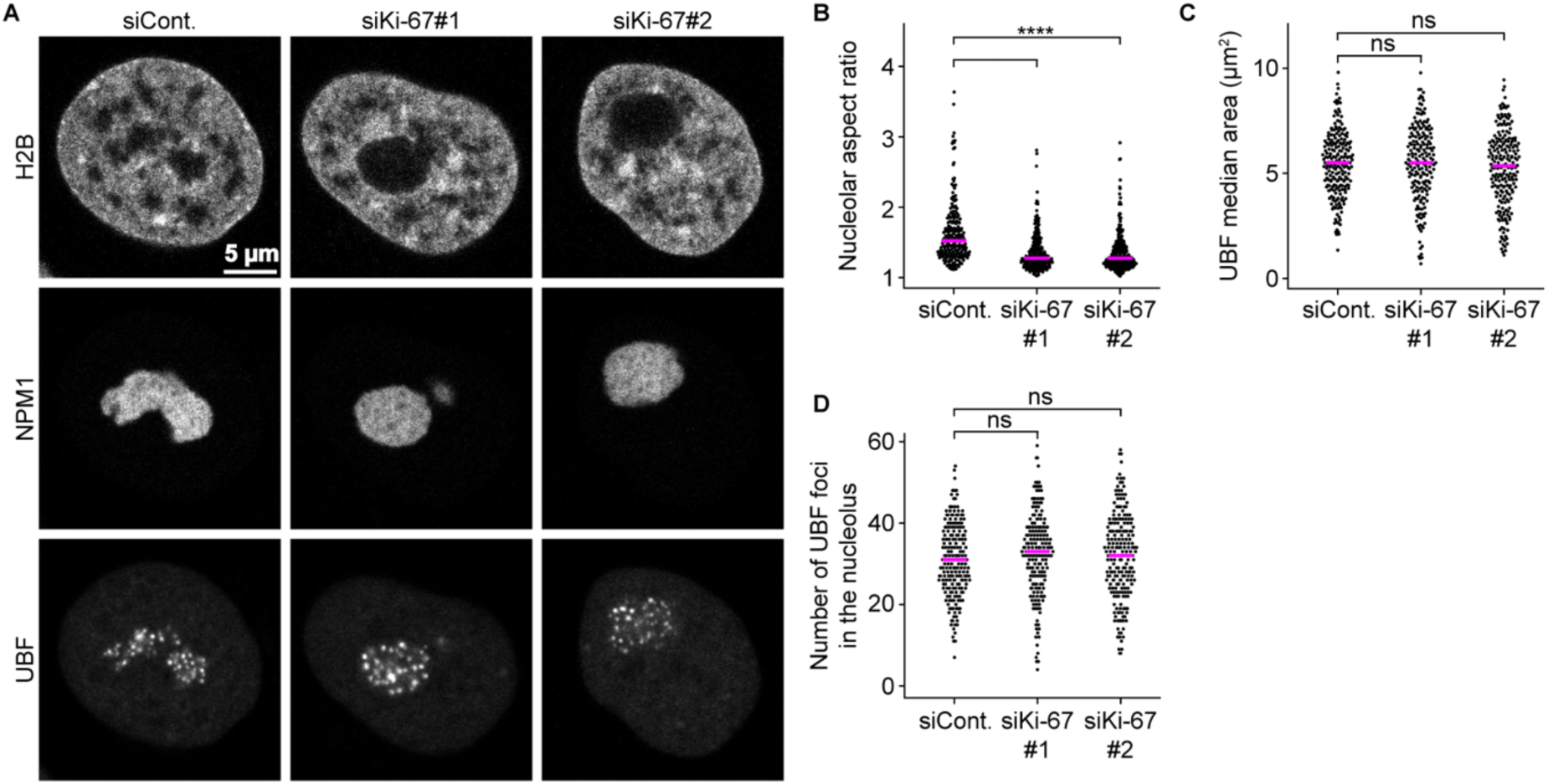
Ki-67 depletion does not disrupt internal nucleolar subcompartments, related to Figure 3. (A) Effects of Ki-67 depletion on nucleolar subcompartments. HeLa cells expressing endogenously tagged Halo-UBF and stably overexpressing SNAP-NPM1 and H2B-mNeongreen were transfected with a non-targeting control siRNA (siCont.) or Ki-67 siRNAs. After 72 h of transfection, Halo-UBF and SNAP-SNAP were labelled with Halo-TMR and SNAP-SiR, respectively. Images were acquired using spinning disk microscopy. (B) Quantification of the nucleolar shape. Nucleoli were segmented based on NPM1 signals, and their aspect ratio was measured. Bars indicate the median. (C, D) Quantification of the size and number of the internal UBF subcompartments based on segmented UBF signals. Median area per the nucleus and the total number of subcompartments were measured. Bars indicate the median. For (B–D), n = 207 nuclei (siCont.), 194 nuclei (siKi-67#1), 214 nuclei (siKi-67#2), 2 experiments. For (B), **** P < 0.0001 with Kruskal–Wallis test followed by Dunn’s test, compared to siControl. For (C, D), ns with ANOVA test.

**Figure S5.**
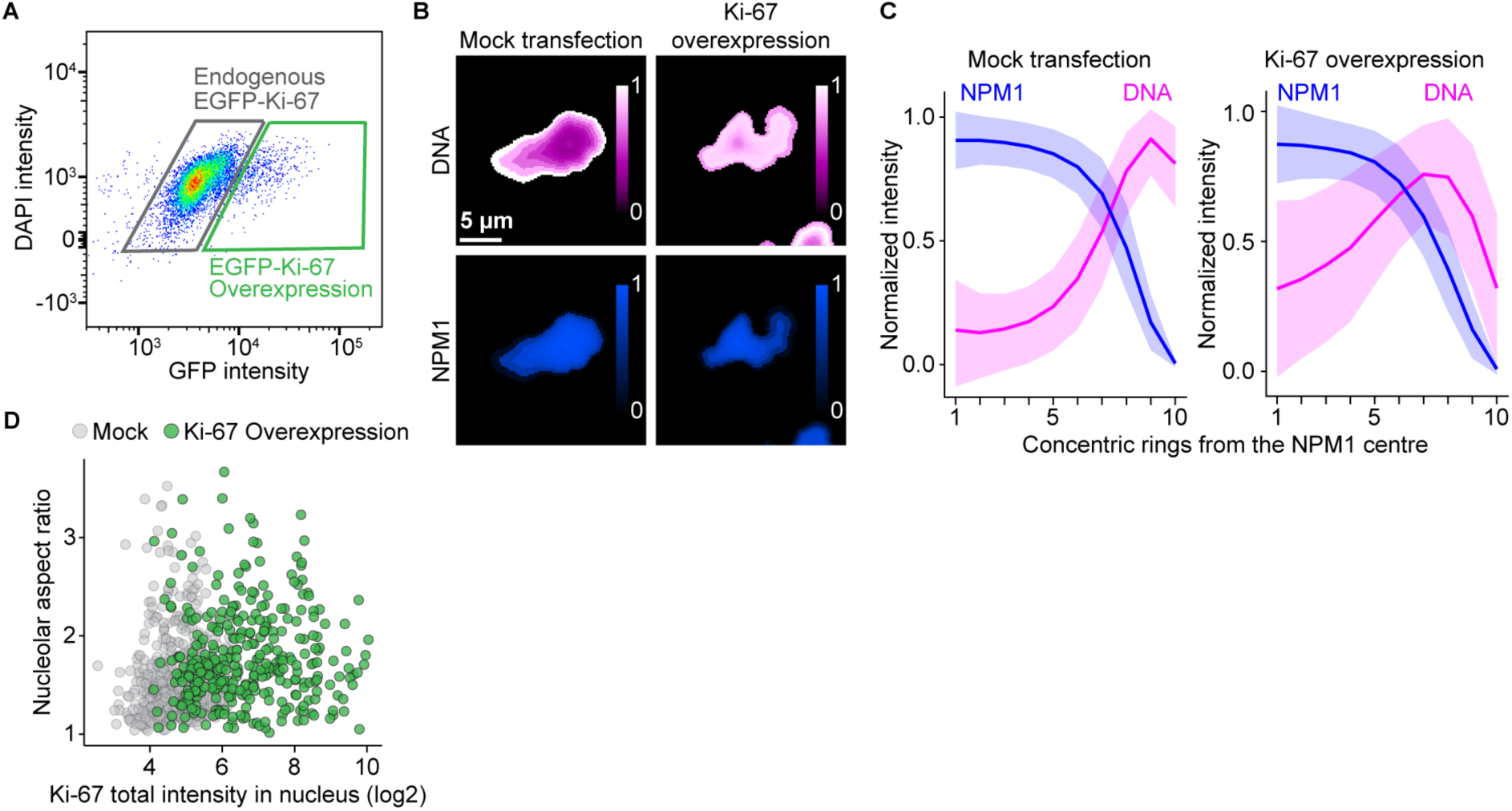
Ki-67 overexpression leads to chromatin incorporation in the nucleolar interior, related to Figure 3. (A) Isolation of Ki-67 overexpressing cells by FACS sorting. Cells expressing endogenous EGFP-Ki-67 were used as a reference (grey gate). Cells expressing higher Ki-67 levels than the reference (green gate) were isolated based on the GFP signal intensities by FACS sorting. (B) Signal distributions of DNA and NPM1 in mock-transfected cells and Ki-67 overexpressing cells. The nucleolus was segmented into 10 concentric rings from the centre of the nucleolus to the edge of the expanded nucleolus. The colour scale represents the relative intensity. (C) Quantification of the signal intensity in concentric rings in mock-transfected cells and Ki-67 overexpressing cells. Mean intensities in each concentric ring were normalised by min-max scaling. Line and shaded areas indicate mean ± SD. (D) Minimal effects on nucleolar aspect ratio upon Ki-67 overexpression. Median nucleolar aspect ratio as a function of the total intensity of EGFP-Ki-67 in the nucleolus reveals no correlation despite resulting in the formation of irregularly shaped nucleoli (Figure 3F). For (H), n = 517 nuclei (mock); n = 355 nuclei (Ki-67 OE), 2 experiments.

**Figure S6.**
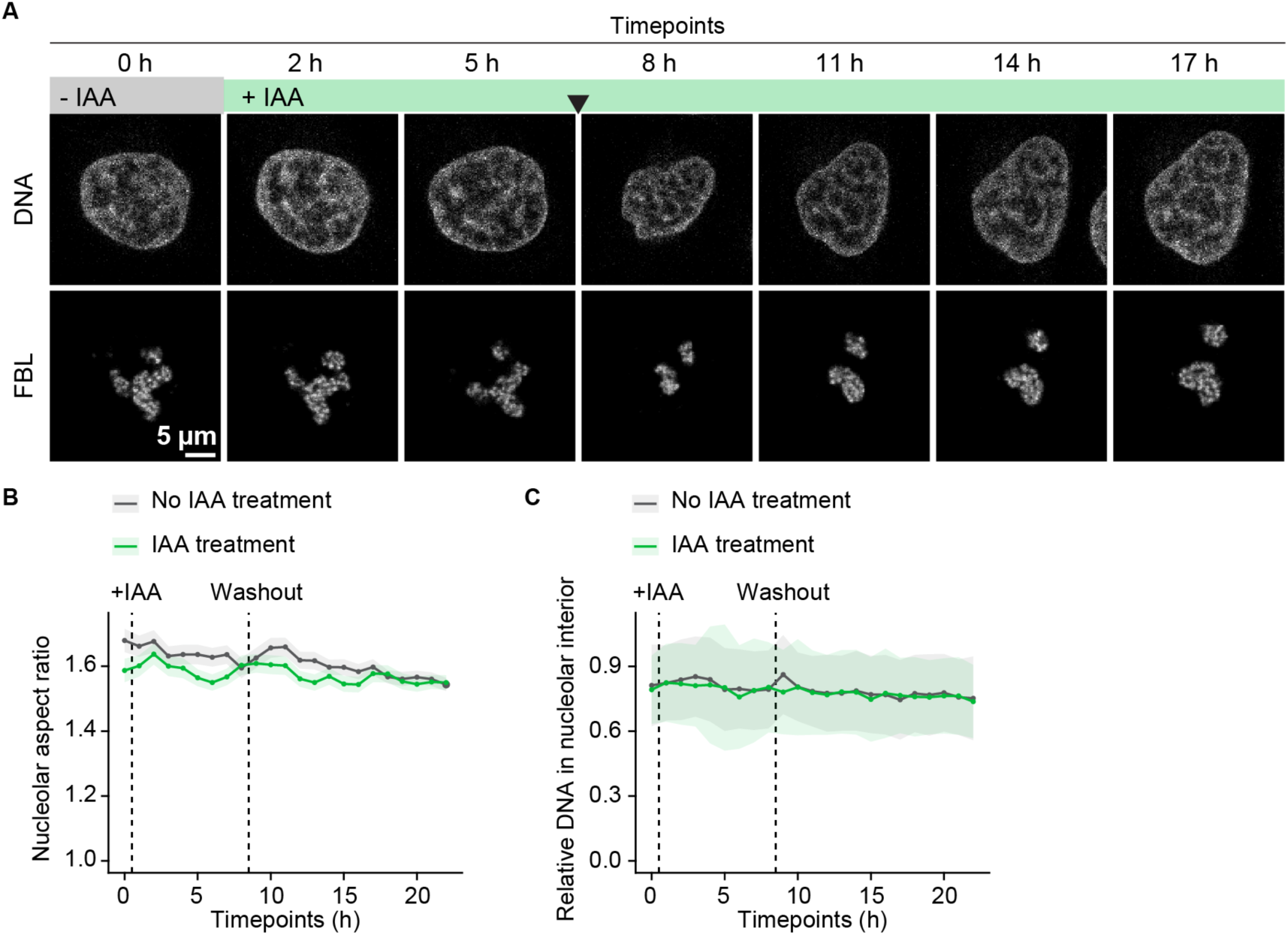
IAA treatment in wild-type Ki-67 cells has no effects on nucleolar shape and chromatin enrichment in the nucleolus, related to Figure 4. (A) Treatment with IAA in control cells expressing wild-type Ki-67 and stably overexpressing FBL-TagRFP. IAA was added 0.5 h after the start of time-lapse imaging. DNA was labelled with SiR-DNA. Single z-slice of a representative example quantified in (B) and (C) is shown. (B, C) Quantification of nucleolar aspect ratio and the DNA enrichment within the nucleolus. The aspect ratio of the nucleolus and the relative DNA mean intensity in the nucleolar interior over the nucleus were measured in non-treated cells (grey) and IAA-treated cells (green). For (B, C), n = 192 to 384 nuclei (No IAA treatment), n = 148 to 363 nuclei (IAA treatment, No washout per time point, 2 experiments.

**Figure S7.**
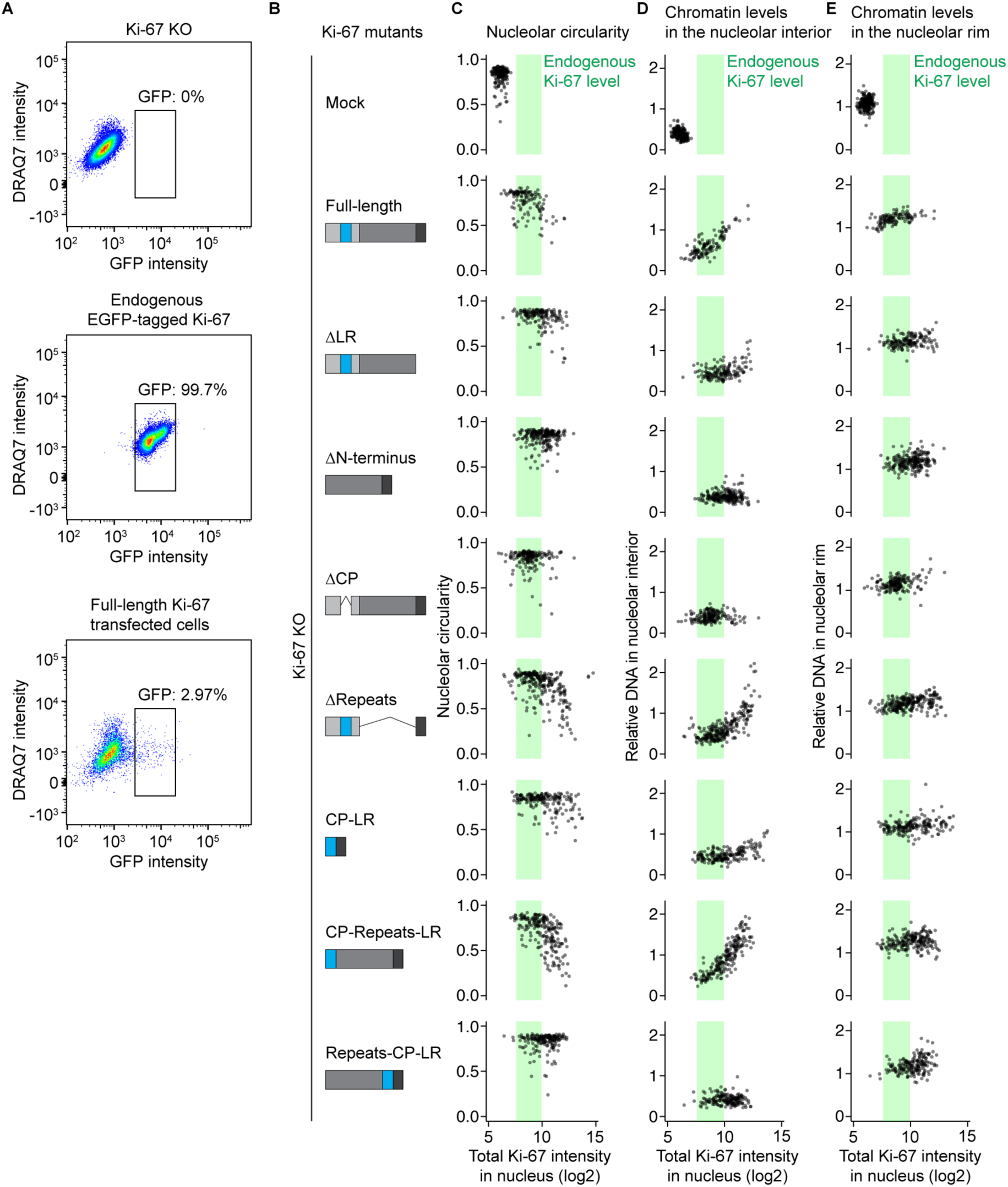
The effect of Ki-67 mutant expression levels on nucleolar shape and chromatin levels within nucleoli and their rim, related to Figure 5. (A) Isolation of Ki-67 mutants with endogenous Ki-67 levels by FACS. Ki-67 KO cells (left) and endogenously EGFP-Ki-67 expressing cells (middle) were used as a reference for FACS sorting. (B) Schematic of Ki-67 domain mutants. (C) Ki-67 expression level-dependent increase in the nucleolar irregularity. Median circularity of nucleoli based on NPM1 segmentation per nucleus is plotted against the total intensity of EGFP-Ki-67 in the nucleus. Green rectangles show the wild-type Ki-67 expression range determined by imaging of endogenously tagged EGFP-Ki-67 cells. (D, E) Ki-67 expression level-dependent chromatin enrichment in the nucleolar interior (D) and rim (E). Relative DNA signal intensities, calculated as described for the H2B signal intensities in (Figure 3D and 3E), are plotted against total EGFP-Ki-67 intensity in the nucleus. Green rectangles show the wild-type Ki-67 expression. For (C–E), n = 220 nuclei (mock), n = 126 nuclei (full-length), n = 162 nuclei (ΔLR), n = 228 nuclei (ΔN-terminus), n = 181 nuclei (ΔCP), n = 128 nuclei (LR), n = 250 nuclei (ΔRepeats), n = 189 nuclei (CP-LR), n = 212 nuclei (CP-Repeats-LR), n = 183 nuclei (Repeats-CP-LR), 3 experiments.

**Figure S8.**
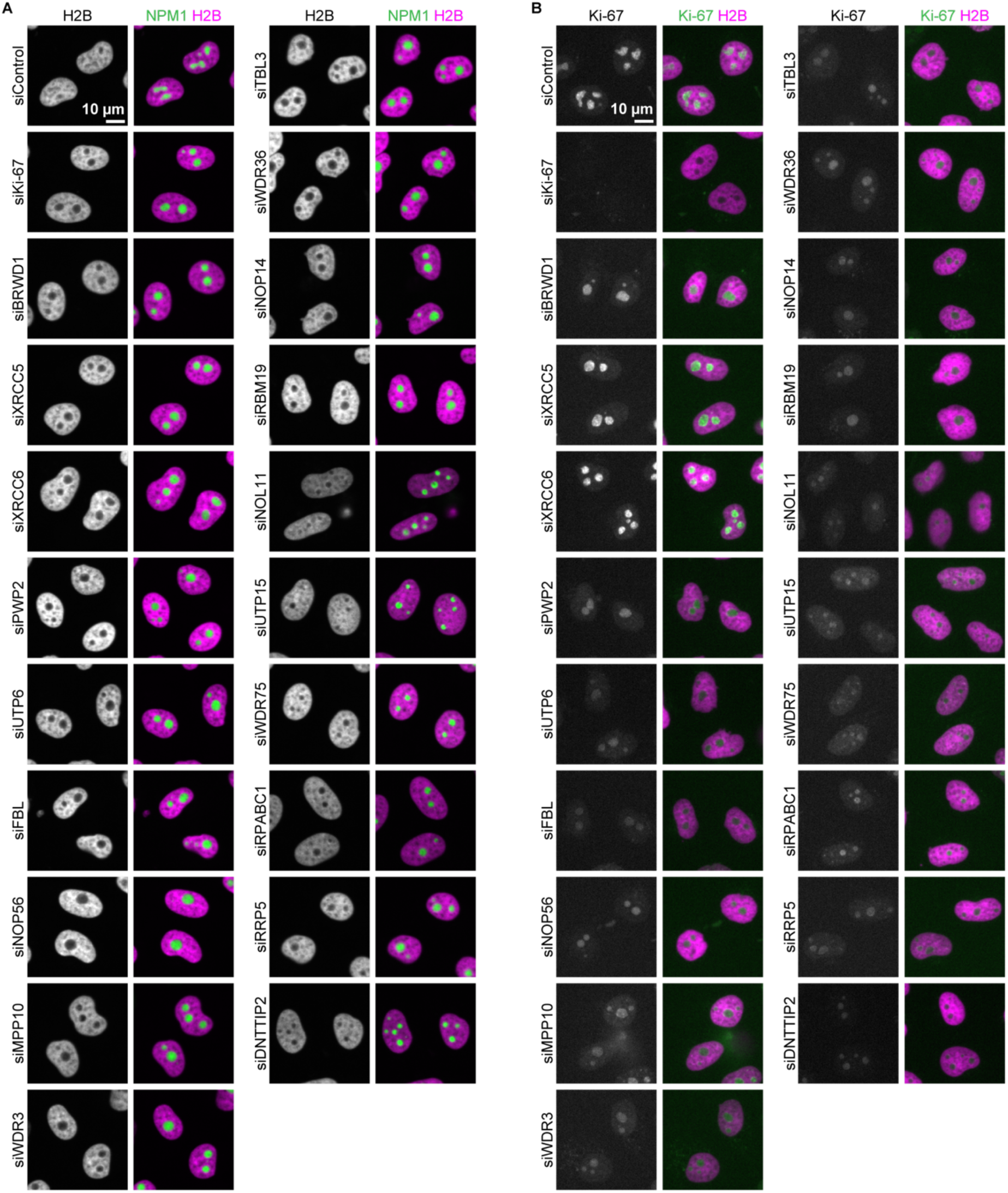
Chromatin enrichment and Ki-67 expression in cells upon siRNA depletion of the top 20 candidate genes inducing nucleolar rounding, related to Figure 7. (A) Live imaging of chromatin enrichment in the nucleolus following depletion of the top 20 candidate proteins. Cells expressing NPM1-EGFP and H2B-mCherry were transfected with a non-targeting control siRNA (siControl) or siRNAs targeting the top 20 candidate genes (Figure 1B). Images were acquired 72 h after siRNA transfection. (B) Live imaging of endogenous EGFP-Ki-67 following depletion of the top 20 candidate proteins. Cells endogenously tagged EGFP-Ki-67 and stably overexpressing SNAP-NPM1 and H2B-mCherry were transfected with a siControl or siRNAs targeting the top 20 candidate genes (Figure 1B). Images were acquired 72 h after siRNA transfection.

