## Supplemental Table 2 and 3 for "Ki-67 shapes the nucleolus by anchoring chromatin via its amphiphilic properties"

**Supplementary Table 2. List of cell lines used in this study.**

| Background | Cell line name | Reference | Lab ID |
|---|---|---|---|
| HeLa Kyoto | Wild-type cells | Originally from S. Narumiya (Kyoto University, Japan) | 1 |
| HeLa Kyoto | NPM1-EGFP, H2B-mCherry | Generated in this study | 36 |
| HeLa Kyoto | siKi-67 no. 2 resistant (homozygous) | Published in [34] | 61 |
| HeLa Kyoto | Ki-67 KO | Published in [34] | 63 |
| HeLa Kyoto | mEGFP-Ki-67 (homozygous) | Published in [34] | 77 |
| HeLa Kyoto | mEGFP-Ki-67 (homozygous), H2B-mcherry | Gift from Daniel W. Gerlich | 80 |
| HeLa Kyoto | FBL-TagRFP (RIEP) | Generated in this study | 161 |
| HeLa Kyoto | mEGFP-miniDegron-Ki-67 (homozygous), OsTir1, FBL-TagRFP (RIEP) | Generated in this study | 163 |
| HeLa Kyoto | Ki67-KO, mTurquoise2-NPM1, FBL-mScarlet (RIEP) | Generated in this study | 349 |
| HeLa Kyoto | mEGFP-Ki-67 (homozygous), FBL-Halo, SNAP-NPM1 | Generated in this study | 391 |
| HeLa Kyoto | mEGFP-Ki-67 (homozygous), H2B-mCherry, SNAP-NPM1 | Generated in this study | 394 |
| HeLa Kyoto | FBL-Halo, SNAP-NPM1 (RIEP) | Generated in this study | 318 |
| HeLa Kyoto | Halo-UBF (homozygous), SNAP-NPM1, H2B-mNeongreen | Generated in this study | 534 |
| HeLa Kyoto | Halo-UBF (homozygous), SNAP-NPM1 | Generated in this study | 567 |
| HeLa Kyoto | Halo-UBF (homozygous), FBL-mEGFP, SNAP-NPM1 | Generated in this study | 395 |

**Supplementary Table 3. List of plasmids used in this study.**

| Name | Comments | Reference | Lab ID |
|---|---|---|---|
| mEGFP-Ki-67 | Full-length Ki-67. | Published in [36] | 343 |
| mEGFP-Ki-67ΔRepeats | Full-length Ki-67 lacking its 16xRepeats (aa 1003 – 2929). | Published in [36] | 607 |
| mEGFP-Ki-67ΔN-term | Full-length Ki-67 lacking its N-terminal region (aa 1 – 1002) | Published in [36] | 665 |
| mEGFP-Ki-67ΔCP | Full-length Ki-67 lacking its charged patch (CP, aa 496 – 681). | Published in [36] | 483 |
| mEGFP-Ki-67ΔLR | Full-length Ki-67 lacking its LR domain (aa 2930 – 3256). | Published in [36] | 726 |
| mEGFP-CP-LR | Charged patch (CP, aa 496 – 681) fused to LR-domain (aa 2930 – 3256). | Generated in this study | 757 |
| mEGFP-CP-Repeats-LR | Charged-Patch (CP, aa 496 – 681) fused to C-terminus of Ki-67 containing 16xRepeats and the LR-domain (aa 1003-3256). | Published in [36] | 818 |
| mEGFP-Repeats-CP-LR | 16xRepeats (aa 1003 – 2929) fused to the charged patch (CP, aa 496 – 681) fused to the LR domain (aa 2930 – 3256). | Generated in this study | 819 |
